# RIF1-ASF1-mediated high-order chromatin structure safeguards genome integrity

**DOI:** 10.1101/2021.11.19.469338

**Authors:** Sumin Feng, Sai Ma, Kejiao Li, Shengxian Gao, Shaokai Ning, Jinfeng Shang, Ruiyuan Guo, Britny Blumenfeld, Itamar Simon, Qing Li, Rong Guo, Dongyi Xu

**Affiliations:** State Key Laboratory of Protein and Plant Gene Research, School of Life Sciences, Peking University, Beijing, China 100871.; Department of Microbiology and Molecular Genetics, Institute of Medical Research Israel-Canada, Faculty of Medicine, The Hebrew University, Jerusalem 91120, Israel

**Keywords:** BRCA1, 53BP1, RIF1, ASF1, ASF1a, Shieldin, NHEJ, HR, resection

## Abstract

The 53BP1-RIF1 pathway antagonizes resection of DNA broken ends and confers PARP inhibitor sensitivity on BRCA1-mutated tumors. However, it is unclear how this pathway suppresses initiation of resection. Here, we identify ASF1 as a partner of RIF1 via an interacting manner similar to its interactions with histone chaperones CAF-1 and HIRA. ASF1 is recruited to distal chromatin flanking DNA breaks by 53BP1-RIF1 and promotes non-homologous end joining (NHEJ) using its histone chaperone activity. Epistasis analysis shows that ASF1 acts in the same NHEJ pathway as RIF1, but via a parallel pathway with the shieldin complex, which suppresses resection after initiation. Moreover, defects in end resection and homologous recombination (HR) in BRCA1- deficient cells are largely suppressed by ASF1 deficiency. Mechanistically, ASF1 compacts adjacent chromatin by heterochromatinization to protect broken DNA ends from BRCA1-mediated resection. Taken together, our findings identified a RIF1-ASF1 histone chaperone complex that promotes changes in high-order chromatin structure to stimulate the NHEJ pathway for DSB repair.

## INTRODUCTION

Double-strand breaks (DSBs) are one of the most cytotoxic DNA lesions and must be effectively and accurately repaired to prevent genomic instability, carcinogenesis, and cell death. To this end, cells must properly choose from two mutually exclusive major DSB repair pathways, homologous recombination (HR) and non-homologous end joining (NHEJ), based on cell cycle position and the nature of the DNA end. DNA end resection, in which broken DNA is converted into 3′-overhang ends suitable for HR, plays a central role in determining the DSB repair pathway choice and is controlled by functional antagonism between the HR-promoting factor BRCA1 and NHEJ-promoting proteins 53BP1 and RIF1 ^1–7^. Mutations in *BRCA1* or *BRCA2* genes cause breast, ovarian, prostate and other cancers, and tumors with such mutations show PARPi hypersensitivity ^8,9^. Unfortunately, PARPi resistance is frequently acquired in patients with advanced cancer. In addition to restoration of BRCA1/2 expression or function by secondary mutations, loss of 53BP1 or its downstream effectors is one major mechanism of PARPi resistance ^10^.

End resection is initiated at proximal chromatin flanking DSBs by BRCA1- promoted CtIP-MRN, and is extended by EXO1 and DNA2/BLM. Recent studies reveal that the shieldin complex (REV7-SHLD1-SHLD2-SHLD3), which is downstream of 53BP1-RIF1, counteracts end resection through CST- and Polα- dependent fill-in ^11–20^. As the shieldin-CST-Polα pathway acts on single-strand DNA (ssDNA) ^13,16,17,20^, this pathway acts like a retrieval system after resection is initiated by mistakes (or in a “trial-and-error” way between HR and NHEJ), or as a restriction system to limit over-resection. In principle, it’s more critical for NHEJ to suppress endonuclease-mediated resection on chromatin flanking DSBs at the initiation step in comparison with the extension step. However, whether and how this function is executed by the 53BP1 pathway remains unclear, although it has long been postulated that the 53BP1-RIF1 complex strengthens the nucleosomal barrier to end-resection nucleases ^16,21,22^.

ASF1 is a histone chaperone conserved from yeast to human cells. Higher eukaryotes contain two paralogs of yeast ASF1, ASF1a and ASF1b, which are distinguishable by their C-terminal tails ^23^. ASF1 functions by transferring H3-H4 heterodimers to the histone chaperone CAF-1 or HIRA for nucleosome assembly ^23^ and contributes to heterochromatin formation ^24–26^. In addition to its role in nucleosome assembly, ASF1 also plays a role in nucleosome disassembly and histone exchange ^27–30^. Here, we find that ASF1 forms a complex with RIF1 in response to DNA damage through a B-domain, which is also responsible for the interactions of CAF-1 and HIRA with ASF1. ASF1 promotes formation of high-order chromatin structure, antagonizes BRCA1-dependent DNA end resection and stimulates NHEJ via its histone chaperone activity. Thus, we identify a RIF1-ASF1 histone chaperone complex that protects broken DNA ends in a parallel pathway with shieldin and confers PARPi sensitivity on BRCA1-deficient cells.

## RESULTS

### The shieldin complex only plays a partial role of RIF1 to promote NHEJ

Previously, we and other groups identified the shieldin complex as a downstream effector of RIF1 to antagonize BRCA1-medicated HR and promote NHEJ ^13–19^. Cell survival experiments using a MTT assay showed that *rif1^-/-^* cells were more sensitive to ICRF193, a topoisomerase II inhibitor that induces DSBs and is specifically toxic to cells deficient in NHEJ ^31–33^, in comparison with SHLD2/FAM35A-deficient cells (see Fig. 1b in reference 13). We confirmed this result using a colony formation assay with sensitivity greater than that of the MTT assay (Supplemental Fig. S1A). PARPis induce one-end DSBs during replication and are toxic to cells deficient in HR proteins such as BRCA1. Disruption of the 53BP1-RIF1 pathway restores HR in BRCA1-deficient cells and thus reduces their PARPi sensitivity ^1–7^. Consistent with the results of the experiments assessing ICRF193 sensitivity, disruption of SHLD2 was not as effective as knockout of RIF1 with regard to rescuing the PARPi (olaparib)-sensitivity of *brca1^-/-^* cells (Supplemental Fig. S1B). Therefore, the shieldin complex only plays a partial role in mediating antagonism of BRCA1 and promotion of NHEJ by RIF1.

**Figure 1.**
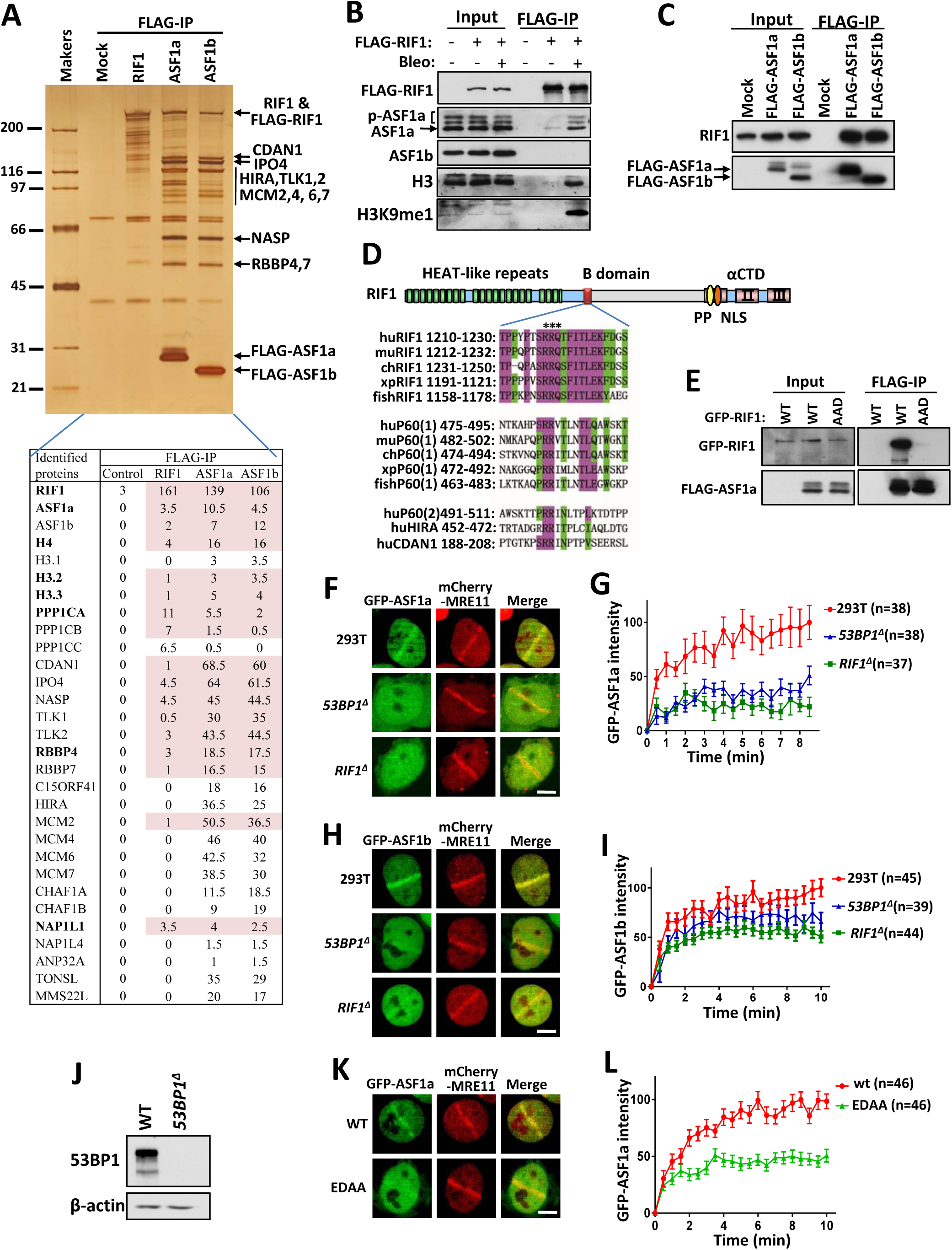
ASF1 forms a complex with RIF1 and is recruited to DSB sites by 53BP1- RIF1. **(A)** A silver-stained SDS-PAGE gel showing the polypeptides that were immunopurified from extracts of HEK293 cells expressing FLAG-tagged RIF1, ASF1a and ASF1b using the anti-FLAG antibody. The major polypeptides on the gel were identified by mass spectrometry. The mean numbers of peptides discovered for each protein from two replicates are listed at the bottom. **(B, C)** Immunoblot showing IP of FLAG-tagged RIF1 (B), ASF1a and ASF1b (C). HEK293 cells expressing FLAG-RIF1 were treated with/without bleomycin (20 μg/mL) for 3 h before harvest. **(D)** Schematic representation of RIF1 and sequence alignment of the B domains of RIF1, CAF-1 P60, HIRA and CDAN1. For proteins with two motifs, both are aligned and designated (1) and (2). hu, *Homo sapiens*; mu, *Mus musculus*; ch, *Gallus gallus*; xp, *Xenopus laevis*; fish, *Danio rerio*. **(E)** Immunoblot showing FLAG-IP from extracts of HEK293 cells expressing FLAG-tagged ASF1a and GFP-RIF1 wild-type (WT) or the AAD mutant. **(F-I)** Recruitment of GFP-ASF1a (F) and GFP-ASF1b (H) to laser-induced DNA damage sites in wild-type, *53BP1^Δ^* or *RIF1^Δ^* HEK293T cells and quantification (G, I). Mean ± SEM is shown for every time point. **(J)** Immunoblots showing protein level of 53BP1 in knockout HEK293T cells. **(K, L)** Recruitment of GFP-tagged wild-type or mutated ASF1a to laser-induced DNA damage sites (K) and quantification (L). Mean ± SEM is shown for every time point. Scale bar, 5 μm.

Additionally, the colony formation assay showed that *shld2^-/-^* cells were more sensitive to etoposide, another topoisomerase II inhibitor that induces DSBs that can be repaired by NHEJ or other pathways ^31,32^, in comparison with *rif1^-/-^* cells (Supplemental Fig. S1A), suggesting that the shieldin complex may have more functions in addition to promoting NHEJ to repair DSBs.

### RIF1 forms a histone chaperone complex with ASF1

To explore the potential parallel pathways of the shieldin complex, we immuno- purified and analyzed the RIF1 complexes from HEK293 cells transiently expressing FLAG-RIF1 with an anti-FLAG antibody. Mass spectrometry analysis revealed that the histone chaperone protein, ASF1a, and H3-H4 were co-immunoprecipitated with RIF1 (Fig. 1A), and immunoblotting confirmed this finding (Fig. 1B). Immunoblotting and mass spectrometry analysis of reciprocal immunoprecipitation showed that RIF1 was present as one major component in the immunoprecipitate of FLAG-ASF1a (Fig. 1A, C), which is consistent with previous interactome analyses ^34–36^. These results demonstrate that RIF1 forms a stable complex with ASF1a-H3-H4.

Moreover, the interaction of ASF1a with RIF1, but not HIRA or CAF-1, was enhanced by DNA damage induced by bleomycin (Fig. 1B; Supplemental Fig. S2A, B), implying that RIF1-ASF1a may play a role in response to DNA damages. ASF1-bound non-nucleosomal H3-H4 heterodimer contains pre-modified H3K9me1, particularly in genotoxic conditions ^37^. Interestingly, H3K9me1 was enriched in the RIF1-ASF1 complex when DNA was damaged (Fig. 1B), implying that RIF1-bound ASF1 tends to provide H3K9me1 upon DNA damages.

ASF1b, a paralog of ASF1a, was not enriched as effectively as ASF1a in the FLAG-RIF1 immunoprecipitate (Fig. 1B), although RIF1 was present as one major component in the FLAG-ASF1b immunoprecipitate (Fig. 1A, C), suggesting that only a subset of RIF1 forms complex with ASF1b.

The downstream histone chaperones CAF-1 p60 and HIRA interact with ASF1 in a mutually exclusive manner via a B-domain motif ^38,39^. Sequence alignment analysis reveals that RIF1 contains a putative B-domain with a high similarity to the domains present in CAF-1 p60, HIRA and CDAN1 (Fig. 1D). Mapping the interacting region revealed that a conserved region (aa1180-1270) of RIF1, which contains the B-domain, was necessary and sufficient for its interaction with ASF1a (Supplemental Fig. S2C-E). Importantly, point mutation in the B-domain of RIF1 (R1217A/R1218A/Q1219D) dramatically reduced its interaction with ASF1a (Fig. 1D, E; Supplemental Fig. S2C). Consistently, mutation of the ASF1a residues (E36A/D37A) critical for binding the B-domain in CAF-1 and HIRA, but not the residue (V94R) for binding H3-H4, disrupted its interaction with RIF1 (Supplemental Fig. S2F, G). Taken together, these results demonstrate that ASF1a binds RIF1 in a manner similar to its interaction with CAF-1 and HIRA, implying that these interactions are mutually exclusive.

### ASF1 is recruited to DSB sites by 53BP1-RIF1

We observed that both GFP-tagged ASF1a and ASF1b were recruited to sites of laser-induced DNA damage (Fig. 1F-I). Recruitment of GFP-ASF1a was dramatically decreased when 53BP1 or RIF1 was disrupted (Fig. 1F, G, J), demonstrating that ASF1a is mainly recruited to DNA damage sites by 53BP1-RIF1. The recruitment of GFP- ASF1b was only modestly reduced in 53BP1- or RIF1-null cells (Fig. 1H, I), suggesting that proteins other than 53BP1-RIF1 also contribute to its recruitment to DSB sites. Consistently, ASF1a mutant EDAA (E36A/D37A), in which residues required for interaction with RIF1 were mutated, showed strongly reduced recruitment to DSB sites (Fig. 1K, L).

### ASF1 colocalizes with 53BP1 and RIF1 at distal chromatin flanking DSBs

53BP1 is excluded from chromatin proximal to DSBs by BRCA1 in the S/G2 phase, resulting in a specific spatial distribution, in which recruitment of BRCA1 into the core of the damage focus is associated with exclusion of 53BP1 to the focal periphery ^40,41^. This distribution of 53BP1 at distal chromatin during HR was deemed to prevent hyper-resection and subsequent single-strand annealing (SSA) to foster its fidelity ^42^. However, due to technical barriers, this spatial distribution has been observed for only a limited number of DSB repair proteins ^40,41^. To examine whether other proteins downstream of 53BP1 are also recruited to chromatin distal to DSB sites, we developed a new method: high-energy-laser-induced recruitment to distal chromatin flanking DSBs (HIRDC). The high-energy-laser-microirradiated regions (1–2-μm- diameter dots) in the cell nucleus are supposed to contain DSB ends, ssDNA and DSB-flanking proximal chromatin, but to largely exclude distal chromatin because of dense DSBs (Fig. 2A). Consistently, ssDNA binding proteins RAD51 and RPA were exclusively recruited in the irradiated dots (Supplemental Fig. S3A), whereas γH2AX localized both inside and outside the dots, although the inside signal was weaker (Supplemental Fig. S3A); BRCA1 and MRE11 were located in regions slightly larger than the dots marked by RPA (Supplemental Fig. S3A), in agreement with reports that BRCA1 and MRE11 are recruited to both ssDNA and proximal chromatin ^43^; Interestingly, both endogenous 53BP1 and GFP-tagged 53BP1 were excluded from the irradiated dots and formed a ring surrounding BRCA1/MRE11 (Fig. 2B, C; Supplemental Fig. S3A-C). The spatial distribution of these proteins was not due to destruction of chromatin by laser microirradiation because staining for H3 and DNA was normal in these regions (Supplemental Fig. S3A). As expected, exclusion of 53BP1 was dose-dependent and impaired when BRCA1 was absent (Supplemental Fig. S3B- D), suggesting that BRCA1 promotes exclusion of 53BP1 from the core region during resection, in agreement with previous studies ^40,41^. Both endogenous and GFP-tagged RIF1 showed re-localization patterns similar to that of 53BP1 (Fig. 2D, E), suggesting that they may work together to prevent hyper-resection during HR in a role similar to their function in promoting NHEJ.

**Figure 2.**
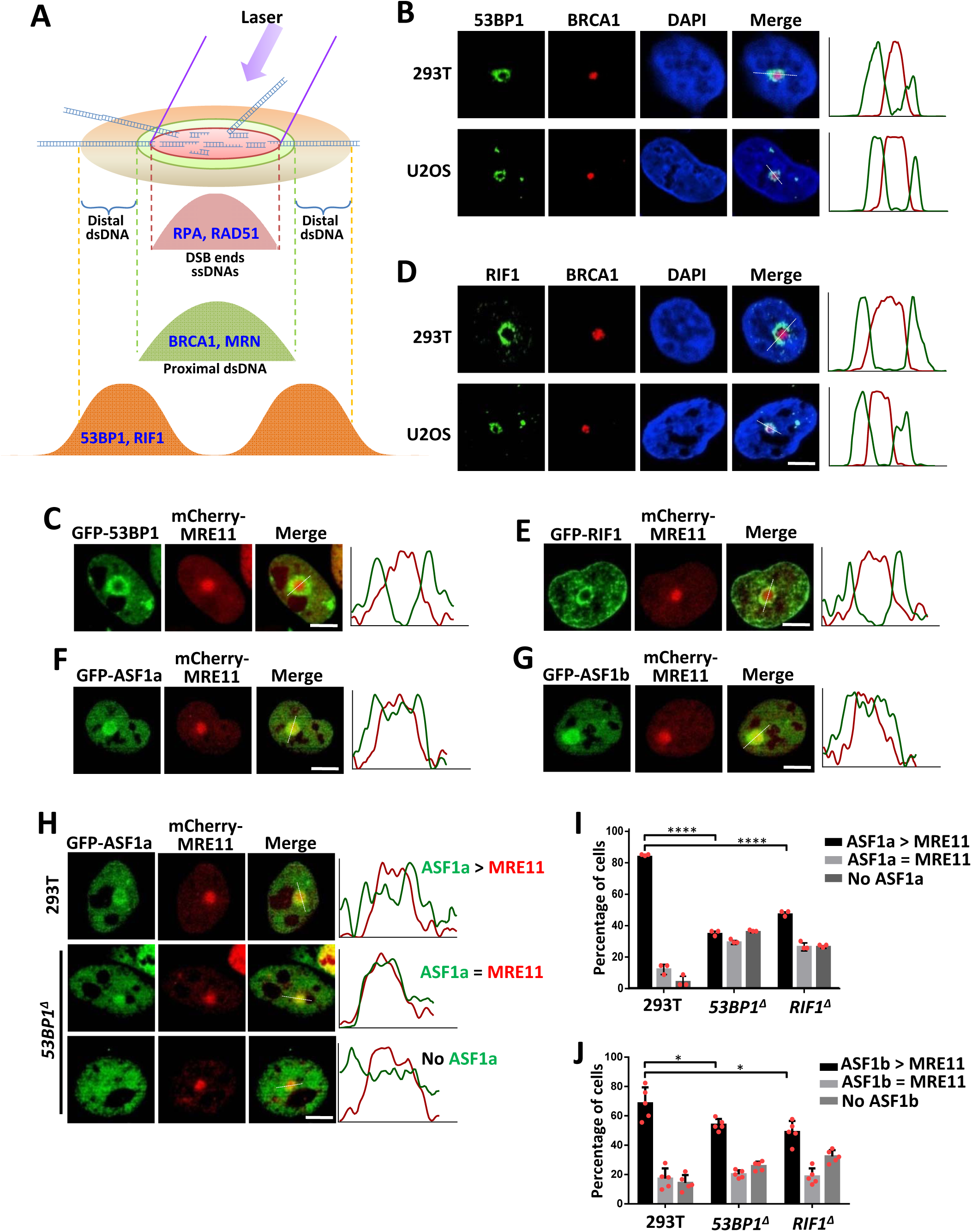
ASF1 is recruited to distal chromatin flanking DSBs by 53BP1-RIF1. **(A)** Schematic representation of the distribution of DNA repair proteins in the HIRDC assay. **(B)** Immunofluorescence showing the distribution of 53BP1 and BRCA1 after micro- irradiation using a high energy laser in the HIRDC assay. The right panel shows the distribution of the red and green signals on the white dashed line as indicated in the left image. **(C)** GFP-53BP1 and mCherry-MRE11 in the HIRDC assay in HEK293T cells. **(D)** Immunofluorescence showing the distribution of RIF1 and BRCA1 in the HIRDC assay. **(E-G)** GFP-tagged RIF1 (E), ASF1a (F) and ASF1b (G) in the HIRDC assay in HEK293T cells. **(H)** GFP-ASF1a in the HIRDC assay in wild-type or *53BP1^Δ^* HEK293T cells. ASF1a > MRE11, the area of accumulated GFP-ASF1a is bigger than that of mCherry-MRE11; ASF1a = MRE11, the area of accumulated GFP-ASF1a is equal with that of mCherry-MRE11; no ASF1a, no accumulated GFP-ASF1a signal. **(I, J)** Quantification of the distributions of GFP-tagged ASF1a (I) and ASF1b (J) in the HIRDC assay in wild-type, *53BP1^Δ^* or *RIF1^Δ^* HEK293T cells. The bars represent the mean ± s.d.; n = 3 (i) and 5 (j) independent experiments; ns, p>0.05; *, p<0.05; ****, p<0.0001; Statistical analysis was performed using the two-tailed *t*-test. Scale bar, 5 μm.

Both GFP-tagged ASF1a and ASF1b were able to re-distribute at distal chromatin flanking DSBs (Fig. 2F, G). In *53BP1^Δ^* and *RIF1^Δ^* cells, these distributions of ASF1a and ASF1b were dramatically and modestly decreased, respectively (Fig. 2H-J), demonstrating that ASF1 acts downstream of 53BP1-RIF1, possibly to limit resection.

Different from 53BP1 and RIF1, ASF1a and ASF1b were also recruited to the core irradiated regions, although to a lesser degree in comparison with the periphery (Fig. 2F, G). Recruitment of ASF1a and ASF1b to the core regions was not significantly affected by disruption of 53BP1 or RIF1 (Fig. 2H-J). These results suggest that ASF1 is also recruited to ssDNA or proximal chromatin by other proteins, consistent with the recent report that ASF1 promotes MMS22L-TONSL-mediated RAD51 loading onto ssDNA during HR ^44^.

### ASF1 promotes NHEJ via the same pathway as RIF1 through its histone chaperone activity

Although HEK293T cells lacking either ASF1a or ASF1b were viable, double knockout cells were not available, possibly due to lethality (Supplemental Fig. S4A). We generated ASF1b heterozygous mutants in *ASF1a*-null cells, which showed more than 50% reduced abundance of ASF1b (Supplemental Fig. S4A). Both *ASF1a^Δ^* and *ASF1b^Δ^* cells were sensitive to ionizing radiation (IR), while *ASF1a^Δ^ASF1b^H^* cells were more sensitive to IR in comparison with the single knockout cells (Supplemental Fig. S4B), suggesting that the two paralogs of ASF1 have overlapping functions in DSB repair.

Chicken DT40 cells have only one paralog of ASF1, ASF1a, which is essential ^45^. By fusing an auxin-inducible degron (AID) at its C-terminus, we generated ASF1a conditional knockout DT40 cells (*asf1a^-/-/AID^*), which showed nearly full loss of the ASF1a protein and proliferation when auxin was present (Supplemental Fig. S4C-E). Even without auxin, *asf1a^-/-/AID^* cells, which retained <10% of the ASF1a protein level of normal cells, were sensitive to etoposide and ICRF193 (Fig. 3A; Supplemental Fig. S4D). In fact, *asf1a^-/-/+^* cells, which retained 33% of the ASF1a protein level of normal cells, also showed modest etoposide and ICRF193 sensitivity (Fig. 3A; Supplemental Fig. S4D). These results suggest that ASF1 is important for DSB repair. In contrast, both *asf1a^-/-/+^* and *asf1a^-/-/AID^* cells were not (or only very weakly at low doses) sensitive to topoisomerase I inhibitor, camptothecin (CPT; Supplemental Fig. S4F), which induces one-end DSBs that depend on the HR pathway for repair ^13,31^, suggesting that ASF1 is almost dispensable for HR in DT40 cells. NHEJ is essential for foreign DNA random integration (approximate 300-fold reduction in *ku70^-/-^* cells compared to wild-type DT40 cells; ^4,13,46^. Random integration in *asf1a^-/-/+^* and *asf1a^-/-/AID^* cells was decreased by 3.1-fold and 6.6-fold, respectively (Fig. 3B). Together, these results demonstrate that ASF1 promotes the NHEJ pathway for DSB repair.

**Figure 3.**
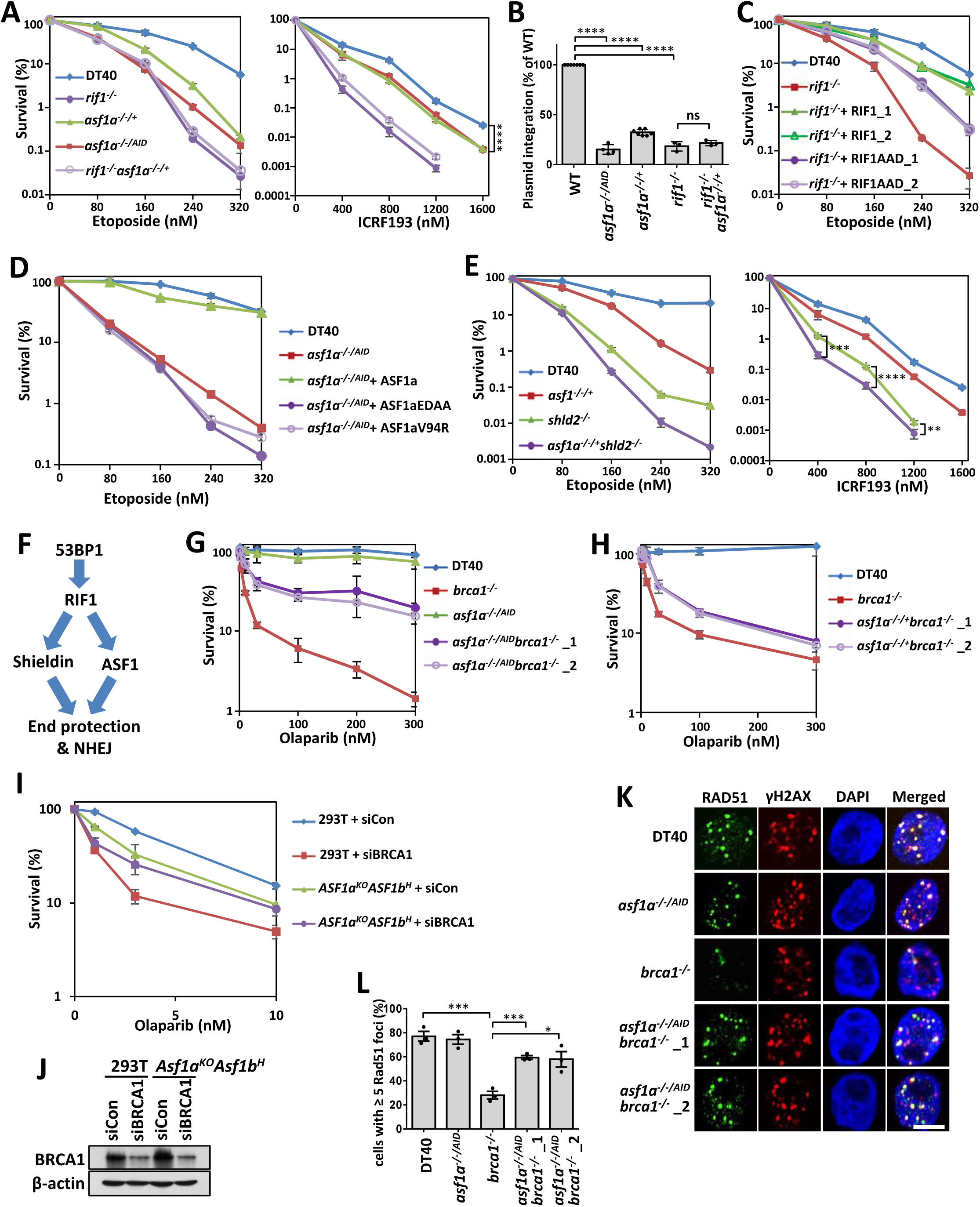
ASF1 suppresses end resection and HR in BRCA1-deficient cells and promotes NHEJ. **(A**, **C**-**E)** Etoposide- or ICRF193-sensitivity assays of various groups of DT40 cells. The mean and s.d. from three independent experiments are shown. **(B)** Random integration assays of various groups of DT40 cells. Data are expressed as the mean number of puromycin resistant colonies ± s.d.; n = 7, 4, 7, 3 and 3 replicates from left to right. **(F)** Cartoon showing the genetic relationships of 53BP1, RIF1, Shieldin and ASF1. **(G, H)** PARPi (Olaparib) sensitivity assay of *brca1^-/-^*, *asf1a^-/-/AID^*, *asf1a^-/-/AID^brca1^-/-^* (G) and *asf1a^-/-/+^brca1^-/-^* (H) DT40 cells. The mean and s.d. from three independent experiments are shown. (**I**) PARPi sensitivity assay for ASF1- and BRCA1-deficient HEK293T cells. The mean and s.d. of the results from two independent experiments are shown. (**J**) Immunoblots showing the knockdown efficiency of BRCA1 in HEK293T cells. **(K, L)** Immunofluorescence (K) and quantification (L) showing that ASF1a blocks end resection in BRCA1-deficient DT40 cells. Cells were treated with 4 Gy X-ray radiation and released 4 h before fixing. The mean and s.d. from three independent experiments are shown. Scale bar, 5 μm. *, p<0.05; **, p<0.01; ***, p<0.001; ****, p<0.0001. Statistical analysis was performed using the two-tailed *t*-test.

We performed genetic interaction analysis to determine whether ASF1 acts via the same NHEJ pathway through which RIF1 functions. The *rif1^-/-^asf1a^-/-/+^* cells did not show more sensitivity to etoposide or ICRF193, or reduction of foreign DNA random integration, in comparison with the corresponding single knockout cells (Fig. 3A, B), demonstrating that ASF1a and RIF1 are epistatic in the NHEJ pathway. Consistently, the interaction between RIF1 and ASF1a was required for promotion of NHEJ: the ASF1a-interacting-defective mutant (AAD) of RIF1 was not as effective as its wild- type form with regard to rescuing the defect of *rif1^-/-^* cells in etoposide resistance (Fig. 3C; Supplemental Fig. S4G); wild-type ASF1a, but not its RIF1-interacting-defective mutant (EDAA), rescued the etoposide sensitivity of *asf1a^-/-/AID^* cells (Fig. 3D; Supplemental Fig. S4H). Thus, ASF1 and RIF1 act in the same pathway to promote NHEJ.

The shieldin complex acts downstream of 53BP1 and RIF1 to protect broken ends and promote NHEJ. Interestingly, *asf1a^-/-/+^shld2^-/-^* cells showed more sensitivity to etoposide and ICRF193 in comparison with *shld2^-/-^* or *asf1a^-/-/+^* cells, demonstrating that ASF1a and the shieldin complex act in two parallel pathways to promote NHEJ downstream of 53BP1-RIF1 (Fig. 3E, F).

Moreover, wild-type ASF1a, but not its histone-binding-defective mutant V94R, rescued the defect of *asf1a^-/-/AID^* cells in resisting etoposide (Fig. 3D; Supplemental Fig. S4H), demonstrating that the histone chaperone activity of ASF1 is required for its function in NHEJ.

### ASF1 protects broken ends and antagonizes HR in BRCA1-deficient cells

We determined whether ASF1 antagonizes the function of BRCA1 in a manner similar to 53BP1-RIF1. As expected, depletion of ASF1 rescued the PARPi-sensitivity of both BRCA1-deficient DT40 and HEK293T cells (Fig. 3G-J). RPA binds to resected ssDNA after end resection and is subsequently replaced by RAD51, which promotes strand invasion and strand exchange during HR ^47^. Reduced foci of both RPA and RAD51 were recovered from BRCA1-deficient cells after ASF1 depletion (Fig. 3K, L; Supplemental Fig. S5A-C), demonstrating that ASF1 opposes HR by limiting resection in BRCA1-deficient cells, similar to 53BP1 and RIF1 ^1–7^.

### 53BP1-RIF1-ASF1 promotes heterochromatinization flanking DSBs

DSBs induce ATM-dependent transcriptional silencing and chromatin condensation at regions flanking damage sites in U2OS-265 cells, which harbor approximately 200 copies of a LacO-TetO array integrated at a single site on chromosome 1p3.6 ^48–50^. Using the same system (Fig. 4A), we confirmed that mCherry-LacR-FokI-caused DSB induced chromatin condensation at the LacO array (Supplemental Fig. S6A-C), while an ATM inhibitor suppressed this condensation (Supplemental Fig. S6A-C). Recently, RIF1 was reported to promote compaction of DSB-flanking chromatin ^51^. Consistently, depletion of 53BP1, RIF1 or ASF1 impaired chromatin condensation at the LacO array after induction of DSBs (Fig. 4B, C; Supplemental Fig. S6D), demonstrating that the 53BP1-RIF1-ASF1 pathway promotes changes in high-order chromatin structure flanking DSBs.

**Figure 4.**
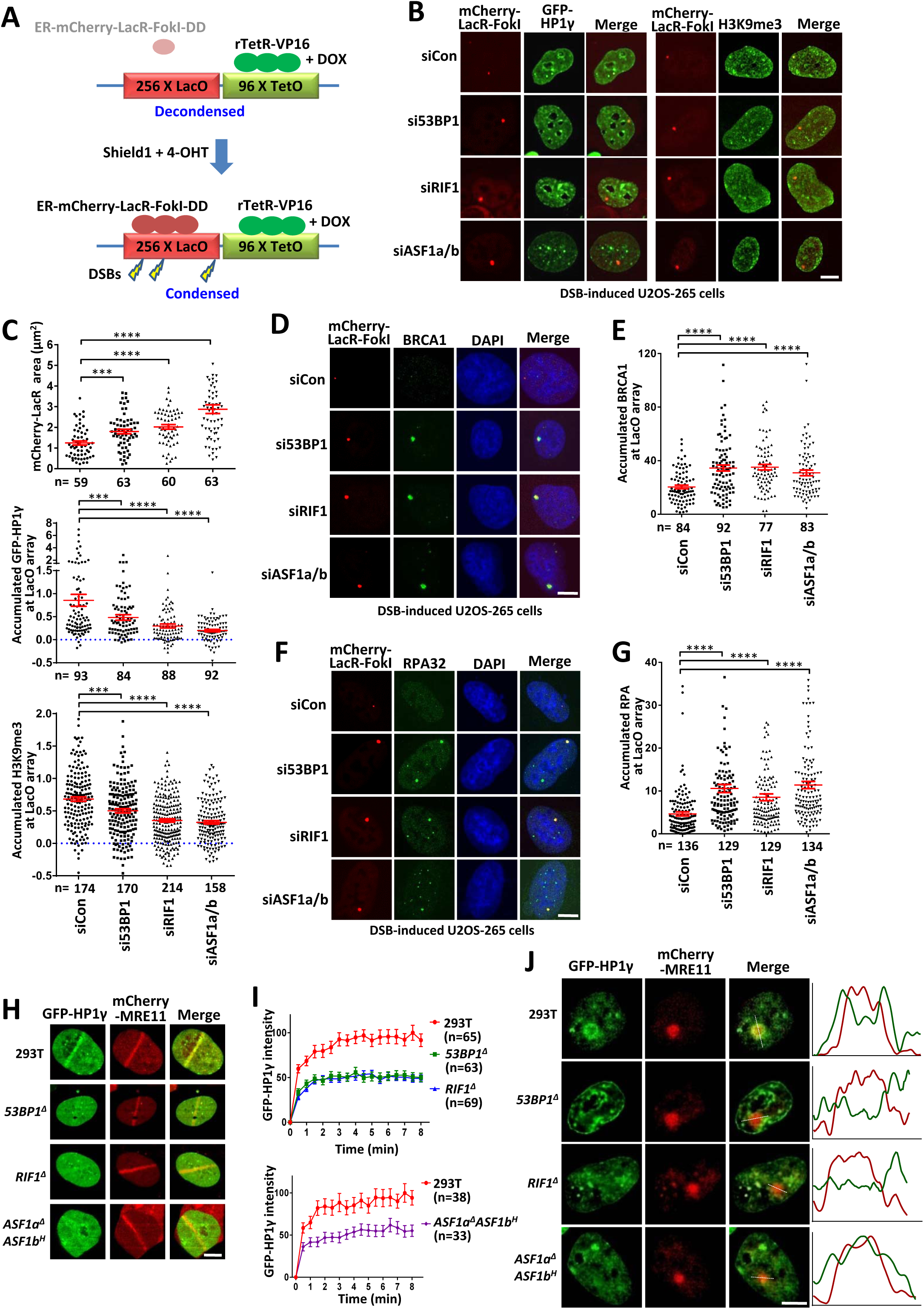
53BP1-RIF1-ASF1 condenses chromatin flanking DSBs. **(A)** Scheme of U2OS-265 DSB reporter cells. Doxycycline (DOX) induces binding of rTetR-VP16 to chromatin and subsequent chromatin decondensation. Shield1 and 4-OHT induce FokI- mediated DSBs within LacO repeats and subsequent chromatin condensation. **(B**, **C)** GFP-HP1γ and H3K9me3 signals (B) in the array and their quantification (C) in 53BP1-, RIF1- or ASF1a/b-depleted U2OS-265 cells after DSB induction. The mCherry-LacR area was also quantified in (C). The mean intensity in the nucleus was set as 0. The error bars represent the SEM. (**D-G**) Immunofluorescence of BRCA1 (D) and RPA32 (F) in the array of DSB-induced USOS-265 cells after depleting 53BP1, RIF1 or ASF1a/b and their quantification (E, G). For RPA32 staining, cells were synchronized in G1 phase as described previously ^92^. The error bars represent the SEM. **(H, I)** Recruitment of GFP-HP1γ (H) to laser-induced DNA damage sites in wild-type, *53BP1^Δ^*, *RIF1^Δ^* or *ASF1a^Δ^ASF1b^H^* HEK293T cells and quantification (I). The mean and SEM values are shown for every time point. **(J)** GFP-HP1γ in the HIRDC assay in wild- type, *53BP1^Δ^*, *RIF1^Δ^* or *ASF1a^Δ^ASF1b^H^* HEK293T cells. Scale bar, 5 μm. ***, p<0.001; ****, p<0.0001. Statistical analysis was performed using the two-tailed *t*-test.

Both RIF1 ^52–57^ and ASF1 ^24–26^ promote heterochromatin assembly in multiple species. Given that RIF1-ASF1 accumulates H3K9me1 (Fig. 1B), a precursor of H3K9me3 for heterochromatin, we thus assessed the impact of RIF1-ASF1 on heterochromatinization at the LacO array upon DNA damage. The signals of heterochromatin markers H3K9me3 and HP1γ were significantly increased at the array after DSB induction (Supplemental Fig. S6A-C), indicating the formation of heterochromatin or heterochromatin-like structures. Interestingly, these signals were decreased when 53BP1, RIF1 or ASF1 was depleted (Fig. 4B, C), demonstrating that these proteins promote chromatin condensation after DNA damage through heterochromatinization.

Chromatin condensation surrounding DSBs via tethering of HP1 or KAP1 suppresses BRCA1-mediated resection ^58^. We determined whether 53BP1-RIF1-ASF1- mediated chromatin condensation at the array suppressed resection. Depletion of 53BP1, RIF1 or ASF1 significantly increased the signals of BRCA1 and RPA32 at the array after DSB induction (Fig. 4D-G), which is consistent with our point that 53BP1- RIF1-ASF1-dependent chromatin condensation antagonizes BRCA1-dependent end resection.

We determined whether 53BP1-RIF1-ASF1-mediated heterochromatinization flanking DSB sites is universal or limited to the artificial sites of the LacO array. GFP- HP1γ was recruited to laser-induced DSB sites and distributed to chromatin distal to DSBs in a manner similar to 53BP1, RIF1 and ASF1 (Fig. 4H-J), suggesting that heterochromatinization occurs at common DSB sites. Recruitment of GFP-HP1γ to laser-induced DSB sites and distal chromatin was significantly decreased when 53BP1, RIF1 or ASF1 was absent (Fig. 4H-J), suggesting that the 53BP1-RIF1-ASF1 pathway promotes universal heterochromatinization flanking DSBs in a manner that is not limited to specific sites.

### RIF1-ASF1 condenses chromatin by promoting heterochromatinization

To assess the molecular mechanism of ASF1 in the formation of higher-order chromatin structure, we monitored changes in the chromatin compaction status of a LacO-TetO array under non-DSB conditions in murine NIH2/4 cells. This array tends to form heterochromatin under normal conditions (Fig. 5A) ^59^. We first decondensed this heterochromatic region by tethering the transcription factor VP16 or the BRCT1 domain of BRCA1, both of which trigger chromatin decompaction ^60,61^, to this array by fusing them with mCherry-LacR (Fig. 5B, C; Supplemental Fig. S7A, B). Interestingly, ASF1a, but not its mutant V94R or EDAA, antagonized VP16 or BRCT1 to re-compact the array when it was fused with GFP-rTetR (Fig. 5B, C; Supplemental Fig. S7A, B). In addition, fusing ASF1a to mCherry-LacR-BRCT1 achieved similar results (Supplemental Fig. S7C, D), which were also validated in human U2OS-265 cells (Fig. 5D, E). Therefore, ASF1a is able to compact opened-chromatin through its histone chaperone activity and its interactions with RIF1 and (or) other histone chaperone, HIRA or CAF-1.

**Figure 5.**
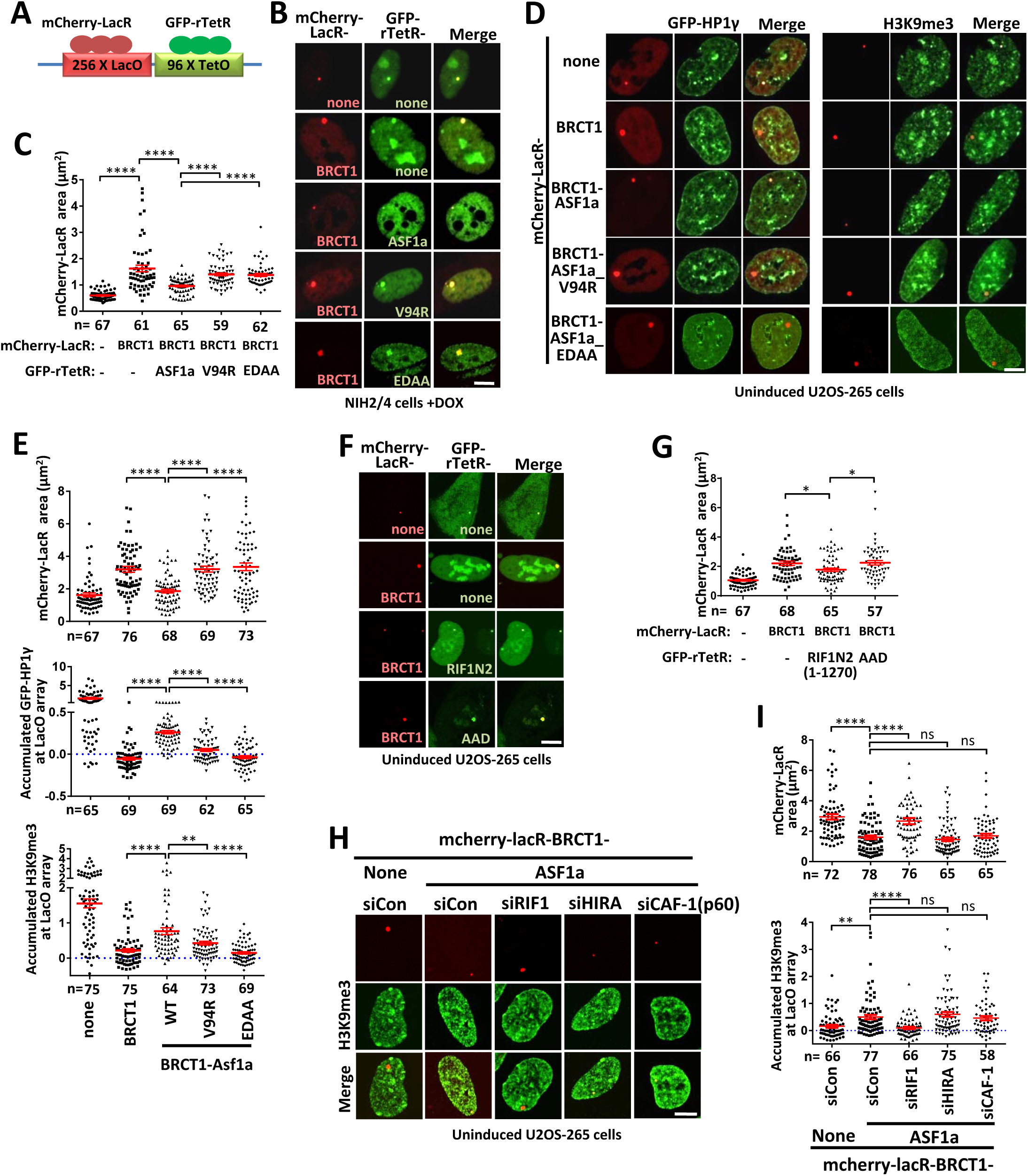
RIF1-ASF1 condenses opened-chromatin by heterochromatinization under non-DSB conditions. **(A)** Scheme of the LacO-TetO array in NIH2/4 cells. **(B, C)** Images (B) and quantifications (C) of NIH2/4 cells expressing GFP-rTetR-fused ASF1a or its mutants (V94R or EDAA), and mCherry-LacR-fused BRCT1 domain of BRCA1 (D, E) with 1 μg/mL DOX. The mCherry-LacR area in the array was quantified. The error bars represent the SEM. **(D)** Left—U2OS-265 cells expressing mCherry- LacR-BRCT1-fused ASF1a or its mutants and GFP-HP1γ. Right—H3K9me3 immunofluorescence in U2OS-265 cells expressing mCherry-LacR-BRCT1-fused ASF1a or its mutants. **(E)** Quantification of the mCherry-LacR area, intensity of GFP-HP1γ and H3K9me3 in the array of experiments as in (D). The mean intensity in the nucleus was set as 0. The error bars represent the SEM. **(F, G)** Images (F) and quantification (G) of U2OS-265 cells expressing GFP-rTetR-fused RIF1N2 or its mutants (AAD), and mCherry-LacR-BRCT1. The error bars represent the SEM. **(H, I)** H3K9me3 signals (H) in the array and their quantification (I) in RIF1-, HIRA- or CAF- 1-depleted U2OS-265 cells expressing mCherry-LacR-BRCT1-ASF1a. The mCherry- LacR area was also quantified in (I). The mean intensity in the nucleus was set as 0. The error bars represent the SEM. Scale bar, 5 μm. ns, p>0.05; *, p<0.05; **, p<0.01; ****, p<0.0001. Statistical analysis was performed using the two-tailed *t*-test.

Tethering the N-terminal region (AA 1-1270) of RIF1, RIF1N2, which contains B domain, but not its mutant AAD, to this array also compacted BRCT1-opened- chromatin under non-DSB conditions (Fig. 5F, G), suggesting that RIF1 is also able to condense chromatin through its interaction with ASF1. Interestingly, ASF1a was no longer able to compact opened-chromatin in RIF1-depleted cells, but not in HIRA- or CAF-1-depleted cells (Fig. 5H, I; Supplemental Fig. S7E). Those results suggest that ASF1a acts in chromatin condensation through forming a complex with RIF1, but not HIRA or CAF-1.

We further examined whether ASF1 compacts chromatin at the LacO-TetO array by promoting heterochromatinization. As expected, tethering of the BRCT1 domain almost fully removed heterochromatin markers H3K9me3 and HP1γ at the array, while tethering of ASF1a, but not its mutant V94R or EDAA, induced a low but statistically significant increase of these signals (Fig. 5D, E). These results demonstrate that ASF1 is able to condensate opened-chromatin by heterochromatinization through its histone chaperon activity and its interaction with RIF1, although its activity level may be low if DSBs are not present. Depletion of RIF1, but not HIRA or CAF-1, suppressed ASF1a-induced-heterochromatinization (Fig. 5H, I). Therefore, the RIF1-ASF1 complex is able to compact chromatin through heterochromatinization in the absence of DNA damages.

### SUV39h1/2 acts downstream of ASF1 to promote heterochromatinization at DSB sites and antagonize BRCA1-dependent HR

SUV39H enzymes catalyze the conversion of H3K9me1 to H3K9me3 and promote the subsequent formation of heterochromatin ^62^. GFP-tagged SUV39h2 was relocalized to laser-induced DNA damage sites and distributed to chromatin distal to DSBs, similar to 53BP1, RIF1 and ASF1 (Fig. 6A-C). And these recruitments of GFP- SUV39h2 were dramatically decreased in 53BP1-, RIF1- or ASF1a/b-deficient cells (Fig. 6A-C), suggesting that SUV39H enzymes may play a role in DSB repair downstream of 53BP1-RIF1-ASF1. The recruitment of SUV39H may be meditated by ASF1-provided H3K9me1 through the affinity between enzyme and its substrate, or (/and) by its interaction with RIF1 as previously described^57^.

**Figure 6.**
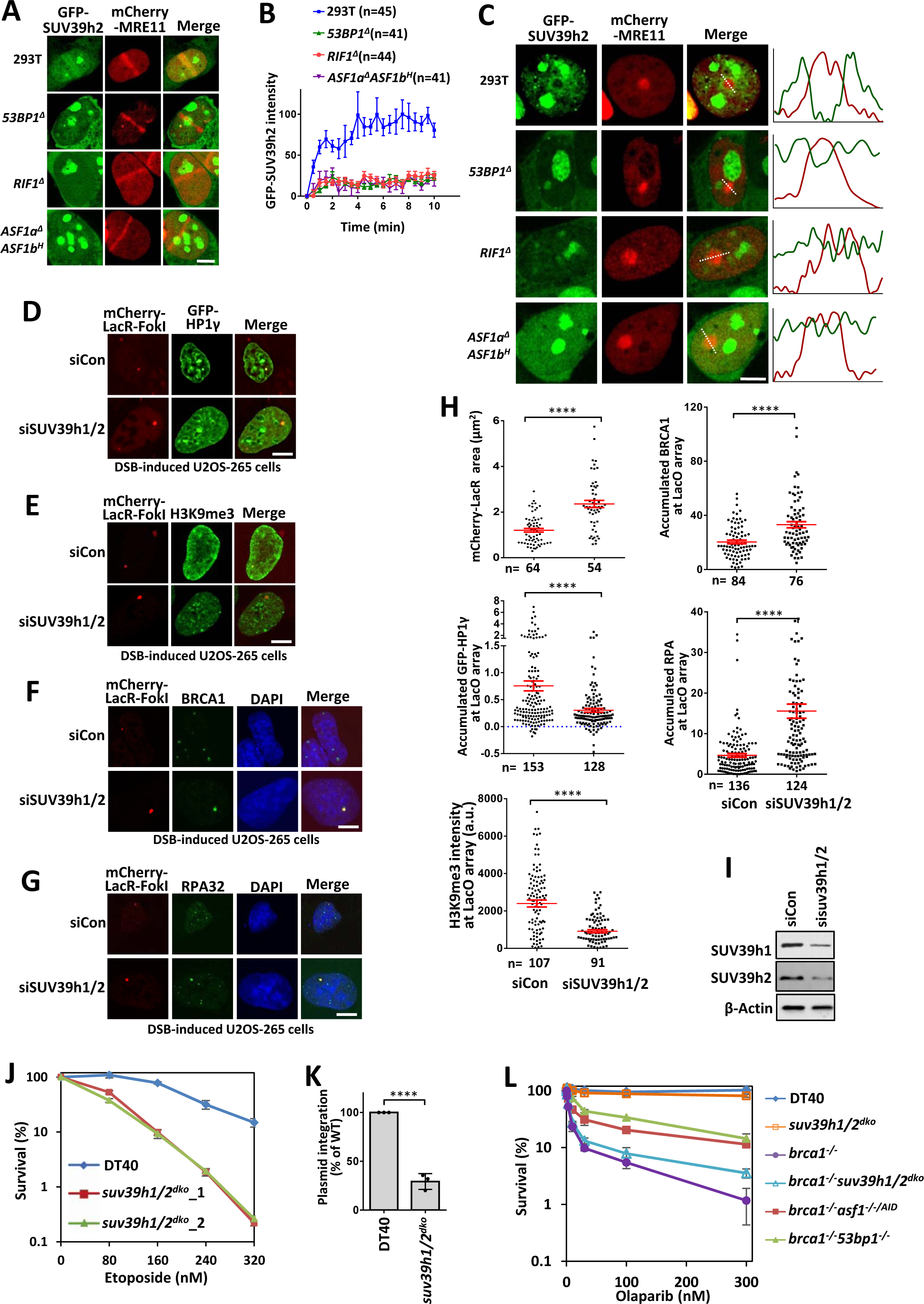
SUV39h1/2 compacts chromatin and antagonizes BRCA1-dependent resection to promote NHEJ. **(A, B)** Recruitment of GFP-SUV39h2 (A) to laser-induced DNA damage sites in wild-type, *53BP1^Δ^*, *RIF1^Δ^* or *ASF1a^Δ^ASF1b^H^* HEK293T cells and quantification (B). The mean and SEM values are shown for every time point. GFP-SUV39h2 in the HIRDC assay in wild-type, *53BP1^Δ^*, *RIF1^Δ^* or *ASF1a^Δ^ASF1b^H^* HEK293T cells. **(D-H)** GFP-HP1γ (D), H3K9me3 (E), BRCA1 (F) and RPA32 (G) signals in the array and their quantification (H) in SUV39h1/2-depleted U2OS-265 cells after DSB induction. The mCherry-LacR area was also quantified in (H). For RPA32 staining, cells were synchronized into G1 phase. Some experiments were carried out together with those in Figure 5B-5G and thus the same controls were used. The error bars represent the SEM. **(I)** Immunoblots showing the knockdown efficiency of SUV39h1/2 in U2OS-265 cells. **(J**, **K)** Etoposide-sensitivity assay (J) and random integration assay (K) of wild-type and SUV39h1/2 double knockout DT40 cells. The mean and s.d. of the results from three independent experiments are shown. **(L)** Olaparib-sensitivity assay of *brca1^-/-^, suv39h1/2^dko^* and *suv39h1/2^dko^brca1^-/-^* DT40 cells. The mean and s.d. of the results from three independent experiments are shown. ****, p<0.0001. Statistical analysis was performed using the two-tailed *t*-test.

As predicted, depletion of SUV39h1 and SUV39h2 prevented chromatin condensation and heterochromatinization, and promoted the recruitment of BRCA1 and RPA32 to the array after DSB induction (Fig. 6D-I), suggesting that SUV39h1/2 promotes heterochromatinization surrounding DSBs and suppresses BRCA1-dependent end resection.

Importantly, disruption of SUV39h1/2 leads to reduced cellular resistance to etoposide and random integration of foreign DNA (Fig. 6J, K; Supplemental Fig. S8A- E), suggesting that SUV39h1/2 promotes NHEJ. Moreover, the absence of SUV39h1/2 rescued PARPi sensitivity of *brca1^-/-^* DT40 cells (Fig. 6L), demonstrating that SUV39H enzymes play a role in antagonizing BRCA1 to suppress HR. The rescue effect of the absence of SUV39h1/2 was weaker than that of deficiency in 53BP1 or ASF1a (Fig. 6L). This difference may have been due to two reasons: one is the activity of a potential parallel pathway to SUV39h1/2 for H3K9 trimethylation; the other is neutralization via the loss of other functions of SUV39h1/2, such as promotion of ATM activity ^63–65^, which is required for PARPi resistance in 53BP1/BRCA1 double-knockout cells ^1^. These findings are in agreement with our conclusion that ASF1-SUV39H1/2-axis-dependent heterochromatinization antagonizes BRCA1 to suppress HR and promote NHEJ.

## DISCUSSION

### RIF1-ASF1 promotes formation of high-order chromatin structure to protect broken ends

It has been long proposed that the 53BP1-RIF1 pathway antagonizes BRCA1- dependent end resection possibly by changing high-order chromatin structure and subsequently controlling nuclease access ^16,22,66^. Here, we identified ASF1 as a partner of RIF1 to protect broken ends through heterochromatinization of the DSB-flanking region (Fig. 7A, B). It is well established that heterochromatin and heterochromatin-like structures represent relatively condensed chromatin that is inaccessible for nucleases. Limiting the accessibility of chromatin neighboring DNA ends not only suppresses excessive resection, but also inhibits its initiation (Fig. 7A, B). This function is quite different from that of shieldin-CST-Polα, which suppresses resection after initiation. Consistently, our data reveal that ASF1 and the shieldin complex act at two parallel pathways to repair DSBs.

**Figure 7.**
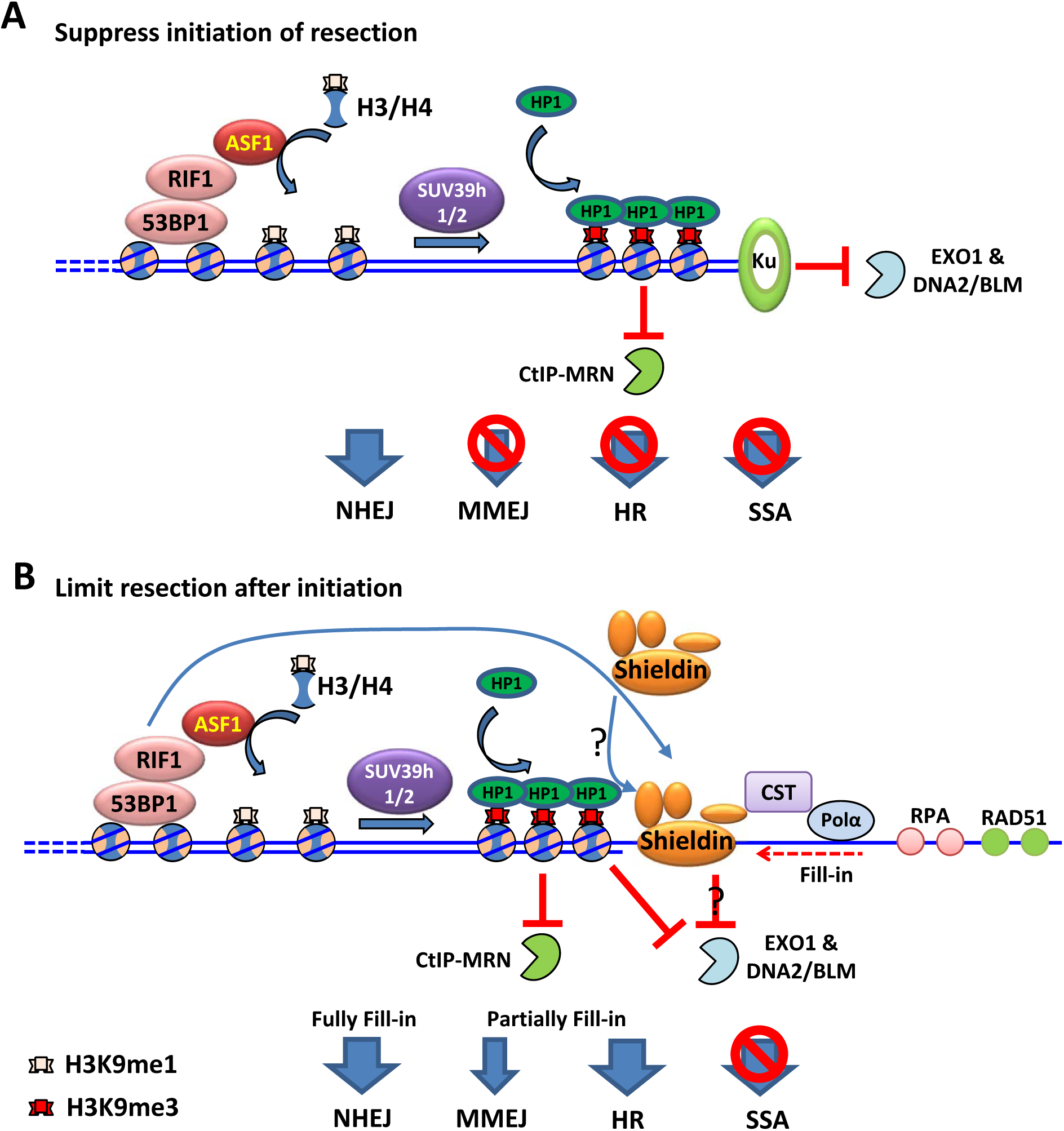
Model of suppression of BRCA1-dependent resection by the RIF1-ASF1 complex. **(A, B)** Models showing the function of RIF1-ASF1 in protection of DNA broken ends. 53BP1-RIF1-ASF1 suppresses the initiation of broken end resection by heterochromatinization, which inhibits microhomology-mediated end joining (MMEJ), HR and SSA (A). After initiation, ASF1 and the shieldin complex cooperate to limit end resection (B). Suppressing resection with full fill-in promotes NHEJ, while limiting resection with partial fill-in promotes MMEJ or HR and inhibits SSA. HP1 might recruit the shieldin complex to DSB sites through direct their interaction, which was shown in several studies ^13,14,18^. HMT, histone methyltransferase.

Assembly of the RIF1-ASF1 complex is dynamically regulated and significantly enhanced in response to DNA damage, although it mimics that of histone chaperone complexes CAF1-ASF1 and HIRA-ASF1. Moreover, upon DSBs, the RIF1-ASF1 complex accumulates H3K9me1, a precursor of H3K9me3 on nucleosomes. It remains to be investigated whether RIF1, like CAF1 and HIRA, has direct histone chaperone activity in the process of H3-H4 deposition, or whether it is only a platform that provides ASF1-H3-H4 to other histone chaperones. Chromatin around DSBs transiently expands (reaching a maximum at about 1.5 min), followed by hypercondensation beyond the predamage baseline level at 20-30 min ^63^. If the expansion accompanied by nucleosome disassembly, RIF1-ASF1 may promote the re- assembly of nucleosome during followed condensation. Otherwise, RIF1-ASF1 may directly promote a nucleosome exchange activity to deposit H3K9me1, which is converted to H3K9me3 for formation of heterochromatin by SUV39H1/2 ^62^. More recently, super-resolution imaging revealed that 53BP1 and RIF1 play a shieldin- independent role in stabilization of compact chromatin topology via an unknown mechanism ^51^. The question of whether ASF1 acts together with 53BP1 and RIF1 in safeguarding chromatin topology around DSBs remains open for future study.

### ASF1 plays multiple functions in DSB repair

It was reported recently that ASF1 promotes MMS22L-TONSL-mediated RAD51 loading onto ssDNA during HR ^44^. Consistently, we detected localization of ASF1 on the ssDNA-region in the HIRDC assay (Fig. 2F, G), and ASF1-defcient cells also showed mild sensitivity to CPT at low dosages (Supplemental Fig. S4F). It’s proposed that NHEJ makes the first attempt to repair DSBs and, if rapid rejoining is not achieved, then the DNA ends are resected and repaired via HR ^67^. The functions of ASF1 in HR and NHEJ may be temporally and spatially separated: ASF1 is initially recruited to the broken ends by 53BP1-RIF1 for NHEJ; if NHEJ repair does not ensure, then after end resection, accumulated ASF1 around DSBs may in turn promote RAD51 loading for HR at ssDNA regions and simultaneously prevent over-resection at adjacent chromatin regions, respectively.

Moreover, ASF1a was reported to interact with MDC1 and promote NHEJ in a manner independent of its histone chaperone activity ^68^. Indeed, unlike RIF1 and MMS22L-TONSL, MDC1 was not detected by mass spectrometry analysis in ASF1a immunoprecipitates from both our (Fig.1A) and other groups ^34–36^, indicating that the interaction between ASF1a and MDC1 may be quite weak or transient. Additionally, unlike that reported in some mammalian cell lines ^68^, histone-binding-defective ASF1a (V94R) could not recover NHEJ activity of *asf1a^-/-/AID^* chicken DT40 cells (Fig. 3D), suggesting that the importance of the histone chaperone activity -dependent and - independent functions of ASF1a in NHEJ is species- or cell- dependent. The question of how the two functions of ASF1a in NHEJ are coordinated in different species and cells remain to be investigated in future.

### Heterochromatin impacts both the efficiency and the balance of HR- and NHEJ-mediated repair of DSBs

Heterochromatic DSBs are repaired more slowly than euchromatic lesions, reflecting that compacted structure of heterochromatin impairs the overall efficiency of DSB repair possibly due to limited accessibility of the sites for repair factors ^69,70^. On the other hand, the balance between HR and NHEJ is strongly influenced by chromatin structure: DSBs that occur in the more open, active chromatin environment of euchromatin are readily repaired by HR, while those occurring in closed, repressive heterochromatin are generally repaired by NHEJ ^58,71,72^. This mechanism may be evolved to avoid genomic instability mediated by HR repair at repetitive DNA elements, which are abundant at heterochromatin ^73,74^. It can be explained that the pre-existing heterochromatin status will facilitate the formation of high-order structure by 53BP1- RIF1-ASF1 after DNA damage, making the balance towards NHEJ repair. Therefore, these studies are in agreement with our model.

Several lines of evidence suggest that heterochromatin marks (HP1, KAP1, SUV39H, H3K9me2, H3K9me3 and macroH2A) and chromatin condensation could take place surrounding DSBs. These marks have both positive ^75–78^ and negative ^58,78,79^ roles in BRCA1-mediated resection and HR. For example, among three HP1 paralogs, HP1α and HP1β stimulates HR, while HP1γ tends to promote NHEJ ^58,78^. Moreover, heterochromatinization is also able to promote ATM-dependent DNA damage response signaling ^63–65^. These contradictory findings can be reconciled to a model in which chromatin condensation may play multiple (even opposite) roles in DSB repair in a spatio-temporally dependent manner (Supplemental Fig. S9). After a transient (about 1.5 min) expansion, chromatin at DSB sites is condensed by heterochromatinization, and then re-relaxes ^63,73,74^. The maintaining time for condensed chromatin is variable from less than 5 min to more than 30 min in different studies possibly due to different chromatin regions, cells and methods ^63–65,80,81^. In a systematic analysis of the kinetics of DNA repair proteins at laser-induced DNA damage sites, HP1 is recruited to damage sites by at least two independent events with a removal halftime of more than 20 min and 60 min, respectively ^80^. This heterochromatinization may be mediated by the 53BP1-RIF1-ASF1-SUV39H axis and protect the broken ends from resection for NHEJ repair at early stage (Supplemental Fig. S9). When rapid repair by NHEJ does not ensure, the heterochromatin mark H3K9me3 then recruits TIP60 to DSBs to hyperacetylate H4 ^82,83^, which leads to chromatin relaxation, promoting BRCA1- dependent end resection and HR ^50^. TIP60 also acetylates and fully activates ATM ^84^, which results in KAP1 phosphorylation and release the KAP-1/HP1/SUV3-9 and CHD3 complexes, promoting the relaxation of heterochromatin ^70,85^. Simultaneously, BRCA1 is recruited by HP1 to promote end resection and HR ^75,77^. Therefore, the temporal and spatial coordination of chromatin dynamics plays a key role in the pathway choice to optimize DSB repair (Supplemental Fig. S9).

Suppression of histone demethylation in isocitrate dehydrogenase 1/2 (IDH1/2)– mutated human malignancies by accumulated 2-hydroxyglutarate causes hyper- trimethylation of H3K9, which confers exquisite sensitivity to PARPi ^86,87^. Our data provide mechanistic insight into how H3K9me3-mediated chromatin compaction causes PARPi sensitivity. Overexpression of epigenetic enzymes such as HADC1/2, SET and SETBD1, which cause chromatin compaction, is widely found in human cancers ^88–90^. Future investigations should assess whether such epigenetic enzymes are biomarkers that could indicate the potential responses of cancers to PARPi therapy.

## Materials and Methods

### Cell culture and transfection

HEK293T, U2OS and NIH2/4 cells were cultured in DMEM medium supplemented with 10% fetal bovine serum (FBS; Invitrogen). HEK293 suspension cells were cultured in SMM 293-TI medium (Sino Biological Inc.) with 1% FBS and 1% glutamine in an incubator with shaking at 140 rpm. The cell lines were obtained from the ATCC and were not among those listed as commonly misidentified by the International Cell Line Authentication Committee. All cell lines were subjected to mycoplasma testing twice per month and found to be negative. The identity of the cell lines was validated by STR profiling (ATCC) and by analysis of chromosome number in metaphase spreads.

DT40 cells were cultured in RPMI 1640 medium with 10% FBS, 2% chicken serum, 10 mM HEPES and 1% penicillin-streptomycin mixture at 39.5 °C (5% CO2). Transfection was performed by electroporation using a Lonza Nucleofector 4D instrument. For selection, growth medium containing G418 (2 mg mL^-1^), puromycin (0.5 μg mL^-1^), blasticidin (25 μg mL^-1^) or histidinol (1 mg mL^-1^) was used.

For knock-down, siRNAs targeting ASF1a (5′-AAGUGAAGAAUACGAUC AAGU-3′), ASF1b (5′-AACAACGAGUACCUCAACCCU-3′), RIF1 (5′-GCAGCU UAUGACUACUAAA-3′), 53BP1 (5′-GAAGGACGGAGUACUAAUA-3′), SUV39h1 (5′-ACCUCUUUGACCUGGACUA-3′), SUV39h2 (5′-UAAUUAUGCUU GUCAUUAGAG-3′), BRCA1 (5′-AAUGCCAAAGUAGCUAAUGUA-3′), CHD1 (5′-GCGGTTTATCAAGAGCTATAA-3′), HIRA (5′- GGAGAUGACAAACUGAUUA-3′), and CAF-1 p60 (5′-AAUCUUGCUCGUCAUACCA-3′) were transfected using RNAi MAX (Invitrogen).

### Antibodies

Antibodies are listed below:

| Antibody Designation | Source or reference | Identifiers | Additional information |
| --- | --- | --- | --- |
| Asf1a(rabbit polyclonal) | Proteintech (China) | 22259-1-AP | WB:1:2000 |
| Asf1b(rabbit polyclonal) | Proteintech (China) | 22258-1-AP | WB:1:2000 |
| Histone3(rabbit polyclonal) | Novus Biologicals<br>(Littleton, USA) | NB500-171 | WB:1:2000 |
| RIF1(rabbit polyclonal) | homemade |  | WB:1:2000 |
| 53BP1(rabbit polyclonal) | Abcam (Cambridge, UK) | ab36823 | WB:1:2000 |
| 53BP1(rabbit polyclonal) | Novus Biologicals<br>(Littleton, USA) | NB100-304 | IF:1:250 |
| BRCA1(rabbit polyclonal) | Millipore (St. Louis, MO, USA) | 07-434 | WB: 1:5000 ;<br>IF:1:1000 |
| BRCA1(mouse monoclonal) | Santa Cruz (Dallas, TX, USA) | sc-6954 | WB:1:100 ;<br>IF:1:40 |
| RAD51(rabbit polyclonal) | Santa Cruz (Dallas, TX, USA) | sc-8349 | IF:1:250 |
| RAD51(rabbit polyclonal) | Abcam (Cambridge, UK) | ab133534 | IF:1:250 |
| γH2AX (mouse monoclonal) | Millipore (St. Louis, MO, USA) | 05-636 | IF:1:5000 |
| RPA32(rabbit polyclonal) | Bethyl (Montgomery,TX, USA) | A300-244A | IF:1:250 |
| Histone H3K9me3(rabbit polyclonal) | Abcam (Cambridge, UK) | ab176916 | IF:1:1000 |
| Histone H3K9me3(rabbit polyclonal) | Abcam (Cambridge, UK) | ab8898 | IF:1:250 |
| BrdU(mouse monoclonal) | Becton Dickinson | 347580 | IF:1:50 |
| MRE11 | Abcam (Cambridge, UK) | ab214 | IF:1:250 |
| β-actin (mouse monoclonal) | MBL (Japan) | M177-3 | WB:1:1000 |

### Immunoprecipitation/MBP-pulldown

For immunoprecipitation (IP), expression plasmids were transfected into HEK293 suspension cells with polyethyleneimine. HEK293 cells were harvested 64 hours after transfection, after which the pellet was lysed with NTEN buffer (20 mM Tris-HCl [pH 7.5], 150 mM NaCl, 10% glycerol, 0.5% NP40, 10 mM NaF, 1 mM phenylmethylsulfonyl fluoride (PMSF), 1 μg mL^-1^ leupeptin, and 1 μg mL^-1^ aprotinin). The lysate was subjected to ultracentrifugation at 440,000 × *g* for 15 minutes, after which the supernatant incubated with anti-FLAG M2 conjugated agarose beads at 4 °C for 4 hours. The beads were washed four times with IP buffer (20 mM Tris-HCl [pH 7.5], 150 mM NaCl, 5 mM MgCl_2_, 10% glycerol, 0.1% NP40, 1 mM DTT and 1 mM PMSF) and incubated with 400 µg mL^-1^ 3× Flag peptide in IP buffer for 1–2 hours. Subsequently, the eluted complexes were analyzed with sodium dodecyl sulfate-polyacrylamide gel electrophoresis (SDS-PAGE) and quality spectroscopy.

For MBP pulldown, 30 μg pDEST26–MBP–RIF1_B or pDEST26–MBP was transfected into a 30-mL suspension of HEK293 cells using polyethyleneimine. After 64 hours, cells were harvested and lysed in 3 mL NTEN buffer. After ultracentrifugation at 440,000 × *g* for 15 minutes at 4 °C, the supernatant was incubated with amylose resin at 4 °C for 4 hours. The beads were washed four times with NTEN buffer and eluted with 50 μL of phosphate buffered saline (PBS) containing 20 mg mL^-1^ of maltose.

### Generation of HEK293T knockout cells

HEK293T knockout cells were generated using the CRISPR/Cas9 genome-editing system. Briefly, pX330 plasmids ^91^ containing the guide sequences (TACAAACATATGCCTTCCTG for ASF1a, GATGAACTCCTGTCCATGGT for ASF1b, ACCTCTGACCAGAGAGCTGC for 53BP1 and GAAGGTAAAGAACCTGCAAC for BRCA1) were transfected into cells. Single colonies were picked after 10–14 days of culture. The isolated single colonies were subjected to western blotting and DNA sequencing to verify protein loss.

RIF1 knockout cells were generated as described previously ^13^.

### Generation of the DT40 knockout strains

The MultiSite Gateway Three-Fragment Vector Construction Kit was used to generate DT40 knockout constructs for the *asf1a* gene. The primer pairs GGAGCTGTGTATCAGGTTGGTATGTTAG/CCCAGCACCTGAGTAGAGACTCT ATG and CCTGAAGAGCAAAGCTCTTATTAGAGGAAG/CAGCCACATCACCCATACCATGAAC were used to amplify the 5′ and 3′ arms from genomic DNA, respectively. The 5′ and 3′ arms were cloned into the pDONR P4-P1R and pDONR P2R-P3 vector, respectively. A knockout construct was generated by attL × attR recombination of the pDONR-5′ arm, pDONR-3′ arm, pDONR-211 resistance gene cassette and pDEST R4- R3 target vector. The C-terminal region of the *asf1a* gene was amplified from genomic DNA using the primer pair CGGGGTACCATGGCAAAGGTTCAGGTGAA/CCGGAATTCTCACATGCAGTCCATGTGGG and cloned into the pAID1.1-C vector. The knockout constructs were linearized before transfection. The *asf1a^-/-/AID^* cells were verified by western blotting. To generate knockout constructs of SUV39h1 and SUV39h2, the 5′ arm and 3′ arm were amplified from chicken genomic DNA using the primers AAGCACAGAGTGGTTGGGTT/AACCCACTTCGGGAGCATTTG (for the SUV39h1 3′ arm), TCCCTCCACCCGCAATAAAC/GTAACCCACTTCCGAGGGTG (for the SUV39h1 5′ arm), GTATGTATTTATCTCATGTGGATTATTTTGAAGGACAAAACAGGAG/CTGGATGAGCTCAGACCATCAGCAGAG (for the SUV39h2 3′ arm) and AGGGCTGGCGAGGAGGTGAG/CTTTAAATACAGAAACAGATAACAGATTTAGCCATGGTTTCAATTC (for the SUV39h2 5′ arm). The first arm was cloned into the pClone007 vector (Beijing Tsingke biotech TSV-007S) using the pClone007 simple vector kit. The second arm and the resistance genes were successively inserted into the pClone007-3′ arm. The knockout construct was linearized before transfection. The gene knockout clones were validated by genomic DNA PCR.

Generation of *RIF1^-/-^*, *SHLD2^-/-^* and *BRCA1^-/-^* DT40 cells was performed as described previously ^4,13^.

### Cell survival assay

For the cell survival assay, 200–400 cells were plated into each well of a 96-well plate with a range of doses of olaparib. After 72 h of incubation, cells were pulsed for 4 h with CellTiter 96 Aqueous One solution reagent (Promega). Cell viability was measured by a luminometer, and each dosage was measured in triplicate. For camptothecin (CPT), a density of 1000–1500 cells per well was used, and the incubation period was 48 hours.

To perform a clone formation assay using DT40 cells, 200–20,000 cells were seeded into each well of a six-well plate filled with 0.7% methylcellulose medium. The plates were treated with the appropriate dose of etoposide or ICRF193. The number of colonies was counted after 7–14 days of incubation at 39.5 °C.

For the clone formation assay with HEK293T cells, 300–20,000 cells were seeded into each well of a six-well plate containing DMEM medium (10% FBS, 1% P/S). The plates were treated with the appropriate dose of olaparib or exposed to the appropriate dose of X-ray radiation. After 10 days of incubation at 37 °C, the number of colonies was counted.

### Random integration assay

DT40 cells were transfected with a linearized pLox-puro plasmid. After 24 hours, 100 cells were plated into 96-well plates to determine the cell plating rate, and one million cells were plated into 96-well plates with 0.5 μg mL^-1^ puromycin for detection of the random integration efficiency. The number of clones was counted after 5–7 days of incubation. The random integration efficiency was normalized by the plating rate.

### Immunofluorescence

Briefly, U2OS or 293T cells were seeded on polylysine-coated overslips before the experiments. After washing with cold PBS, the cells were pre-extracted with 0.5% TritonX-100 in CSK buffer (20 mM HEPES [pH 7.0], 100 mM NaCl, 300 mM sucrose, and 3 mM MgCl2) for 10 min at 4 °C. Next, the cells were washed three times with PBS and fixed with 3% paraformaldehyde for 10 min at room temperature. The cells were permeabilized for 10 min with 0.5% TritonX-100 in CSK buffer, washed three times with 0.05% Tween-20 in PBS and blocked with 5% BSA for 15 min. Next, the cells were incubated with the primary antibodies for 90 min. After washing, the cells were incubated with the secondary antibodies diluted in 1% BSA/PBS for 30 min. After three washes, the cells were mounted with ProLong Gold antifade reagent with DAPI (Invitrogen). Images were acquired with an ANDOR Dragonfly system on a Leica DMI8 microscope with a 100× oil immersion objective.

### Laser-induced foci and HIRDC

U2OS or HEK293 cells expressing GFP- or/and mCherry-fused proteins were cultured at 37 °C in DMEM medium containing 10% FBS. During microirradiation and imaging, the cells were maintained in a temperature-controlled container with 5% CO2 in glass-bottom dishes (NEST Biotechnology). The laser system (Micro-Point Laser Illumination and Ablation System, ANDOR) was directly coupled to a Leica DMI8 microscope with a 100× oil immersion objective. Images were acquired with ANDOR IQ3 software through an ANDOR IXON camera with an ANDOR Dragonfly system.

In order to obtain images of unperturbed cells, time-lapse image acquisition was begun before laser microirradiation. Microirradiation was performed on an indicated line in the nuclei of cultured cells via micro-point laser illumination (65% output) at the time of the second image. Images were collected every 30 seconds for 10 minutes and analyzed with ImageJ software (NIH). Recruitment was measured by determining the mean fluorescence intensity within the damage region and normalizing this value to the mean fluorescent intensity of the unirradiated nucleus. For each cell, a separate region was measured for background subtraction. The relative fluorescence intensity was calculated by the following formula: RFI(t) = (I_t_ - I_b_) / (I_nu_-I_b_), where *I_t_* is the mean fluorescence intensity of the microirradiated region, *I_b_* is the mean fluorescence intensity of the background, and *I_nu_* is the mean fluorescence intensity of the unirradiated nucleus.

For the HIRDC assay, microirradiation was carried out within a dot (approximately 1 μm in diameter) in the nucleus with a fill-in program. The micro-point laser illumination output was set at 65% unless indicated.

### Imaging quantification in U2OS or NIH2/4 cells

U2OS-265 or NIH2/4 cells expressing mCherry-LacR or GFP-rTetR were cultured at 37 °C in DMEM medium containing 10% FBS. Images were acquired with Imaris software (Bitplane) through an ANDOR IXON camera with an ANDOR Dragonfly system. The area of the array was analyzed using Imaris (Bitplane). Cells were chosen at random for the analysis of the intensity of the indicated signals. The region of the array was defined by the mCherry-LacR signal. The accumulated signals were measured by determining the mean fluorescent intensity of the array region and normalizing this value to the mean fluorescent intensity of the entire nucleus. For each cell, a separate region was measured for background subtraction. The accumulated signal was calculated by the following formula: As = (I_array_ - I_b_) / (I_nu_-I_b_), where *I_array_* is the mean fluorescence intensity of the array region, *I_b_* is the mean fluorescence intensity of the background, and *I_nu_* is the mean fluorescence intensity of the nucleus. All intensity analysis was performed on unprocessed images using ImageJ software.

### Statistics and Reproducibility

All immunoblots were performed at least three times unless otherwise noted in the legend. GraphPad Prism 6 and Excel 2013 were used for statistical analysis. Statistical significance was assessed using the two-tailed Student’s *t*-test. The data were normally distributed and the variance between groups being statistically compared was similar. No statistical methods or criteria were used to estimate sample size or to include/exclude samples. All image analyses were carried out in a double-blind approach.

## Acknowledgments

We thank Weidong Wang for his advice and revisions to the manuscript. We thank the Imaging Core and Mass-spectrum Core at the National Center for Protein Sciences at Peking University for imaging and mass-spectrum analysis. NIH2/4 cells were a gift from Tom Misteli. U2OS-265 cells were a gift from Daniel Durocher under a permission of Roger A. Greenberg. This work was supported in part by Beijing Outstanding Young Scientist Program (BJJWZYJH01201910001005) to L.Q. and the National Natural Science Foundation of China (81672773 and 31661143040) to D.X.

## Author Contributions

S.F., S.M., K.L., S.G., S.N, J.S. and R.G. carried out experiments. I.S., Q.L., R.G. and D.X. designed experiments and interpreted the results. S.F., S.M., K.L. and D.X. wrote the manuscript.

---

**Supplemental Figure S1.**
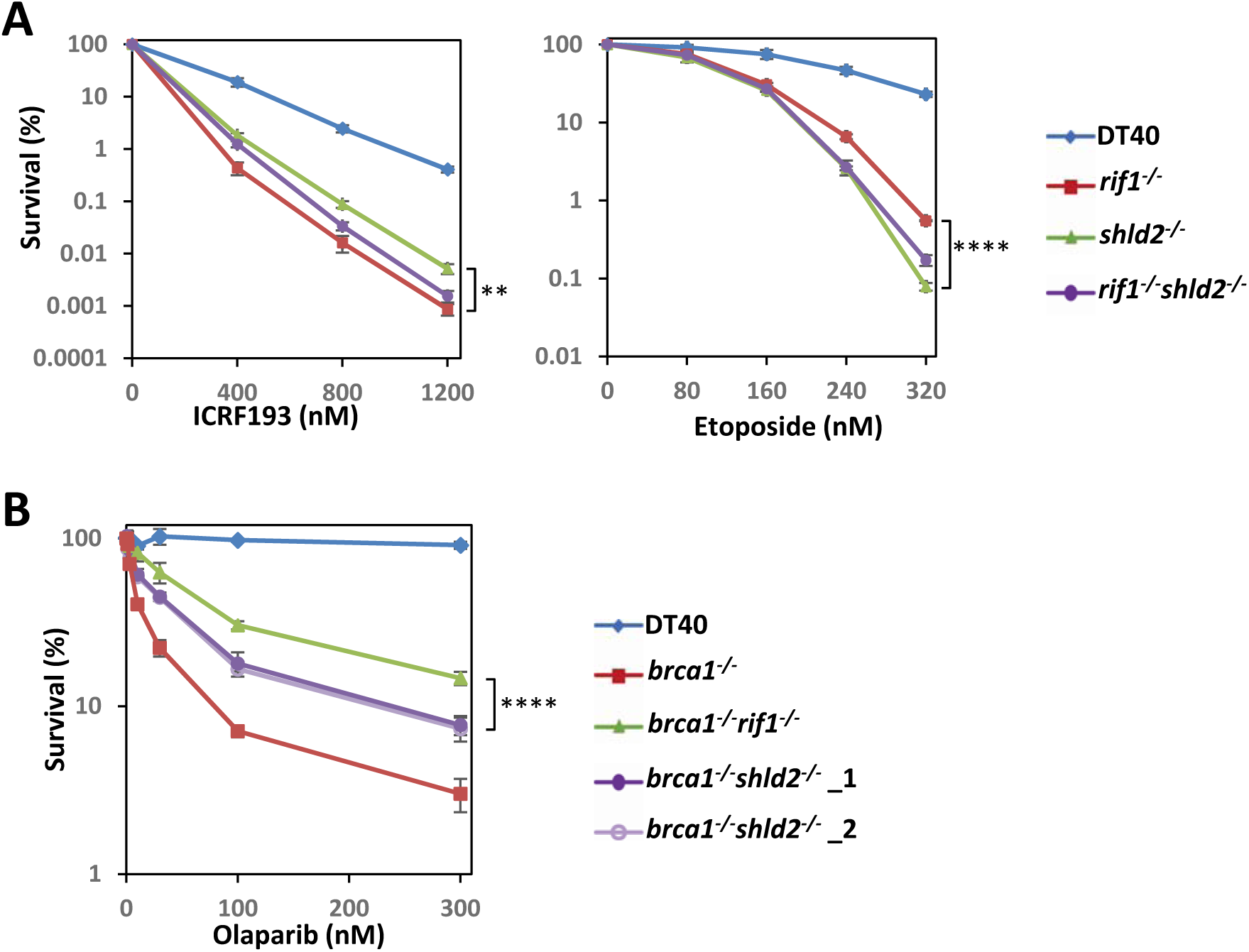
SHLD2 is not as effective as RIF1 to promote NHEJ and suppress resection in BRCA1-deficient cells. (**A**) Etoposide- or ICRF193-sensitivity of various groups of DT40 cells using colony formation assay. The mean and s.d. from three independent experiments are shown. (**B**) The graphic shows the PARPi (olaparib) -sensitivity of BRCA1-deficient DT40 cells when RIF1 or SHLD2 was knockout. Two *brca1^-/-^shld2^-/-^* clones were used. The mean and s.d. from three independent experiments are shown.

**Supplemental Figure S2.**
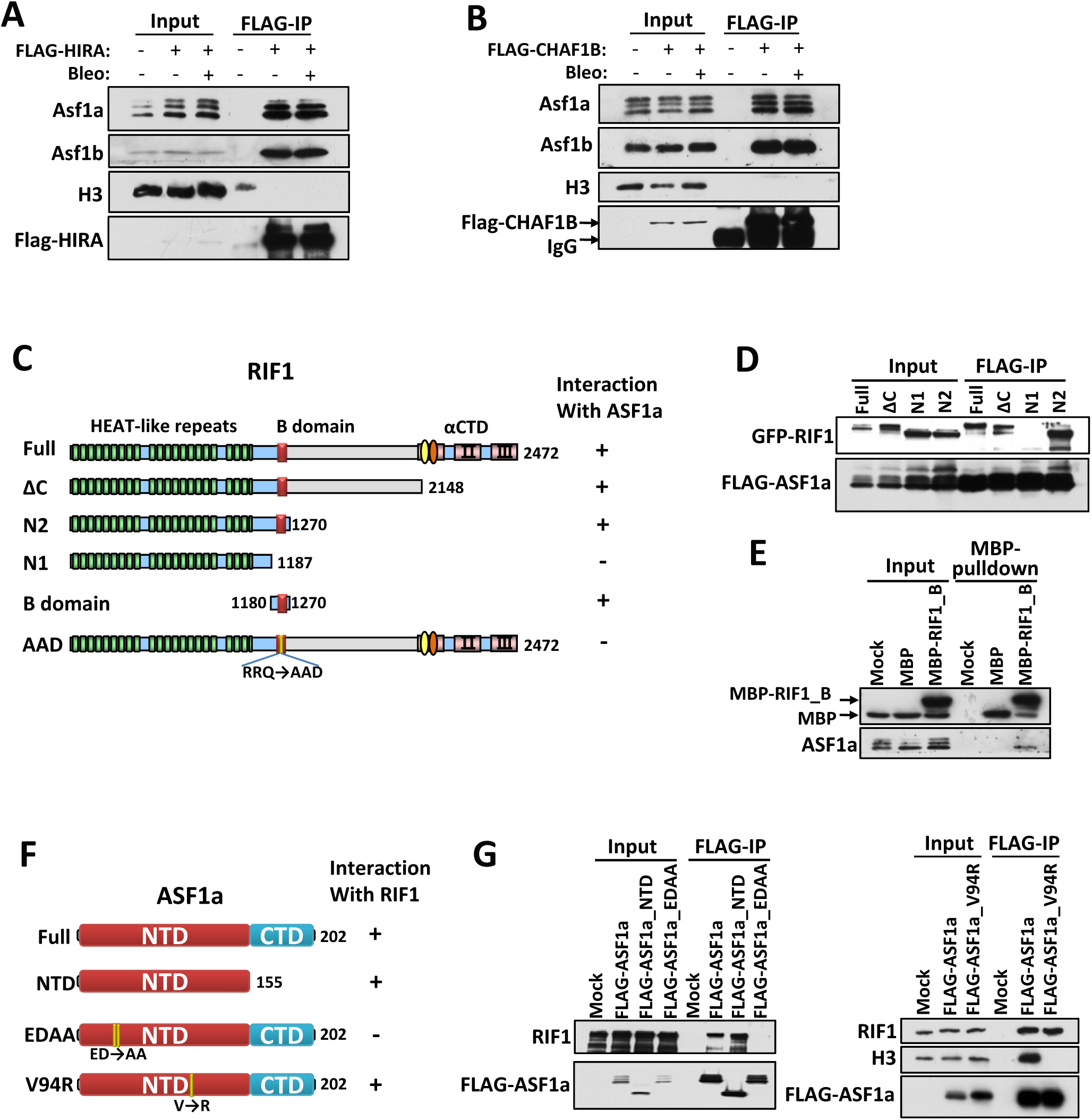
Mapping the interacting regions of RIF1 and ASF1a. (A, B) Immunoblot showing IP of FLAG-tagged HIRA (A) and CAF-1 p60 (CHAF1B) (B). Cells were treated with/without bleomycin (20 μg/mL) for 3 h before harvest. (**C**) Schematic representations of different RIF1 mutants (left) and their ability to coimmunoprecipitate with ASF1a (right). (**D**, **E**) Immunoprecipitation (D) and MBP-pulldown (E) to assess whether the various deletion mutants of RIF1 co-purified with ASF1a. (**F**) Schematic representations of different ASF1a mutants (left) and their ability to coimmunoprecipitate with RIF1 (right). EDAA, E36A/D37A. (**G**) Immunoprecipitation to assess whether the various mutants of ASF1a co-purified with RIF1.

**Supplemental Figure S3.**
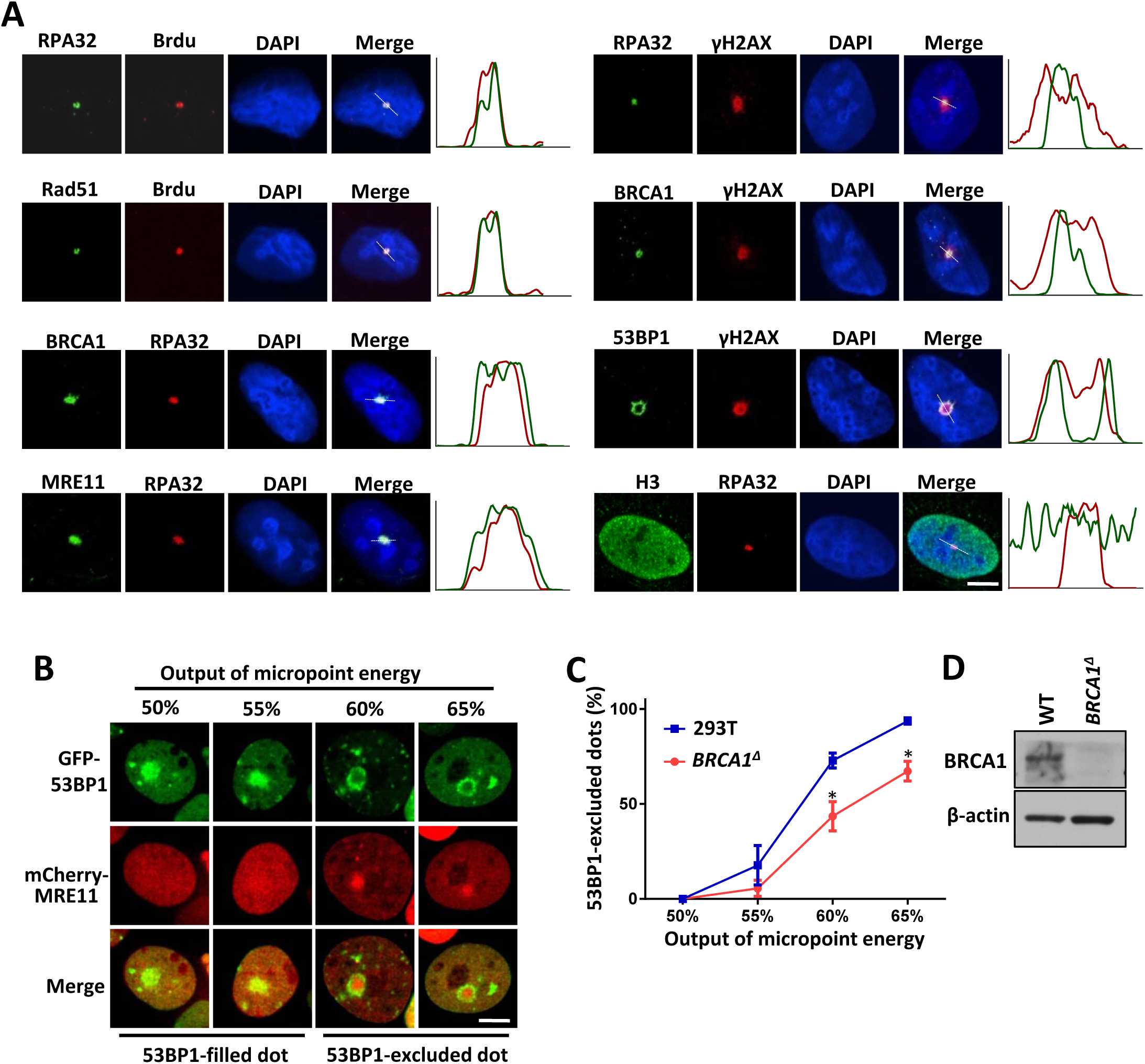
Exclusion of 53BP1 from the core irradiated region is dose-dependent and promoted by BRCA1. (**A**) Immunofluorescence showing the distribution of multiple proteins in the HIRDC assay. For ssDNA staining, cells were incubated with BrdU for 24 hours before micro-irradiation. The right panels for every image show the intensity distribution of the red and green signals on the white dashed line indicated in the image. (**B**) GFP-53BP1 in the HIRDC assay with various doses of laser energy in HEK293T cells. (**C**) Quantification of the distribution pattern of GFP-53BP1 in the HIRDC assay in wild-type or *BRCA1^Δ^* HEK293T cells. Two patterns of GFP-53BP1 distribution were defined as in (B). Scale bar, 5 μm. *, p<0.05; Statistical analysis was performed using the two-tailed *t*-test. **(D)** Immunoblots showing protein level of BRCA1 in knockout cells.

**Supplemental Figure S4.**
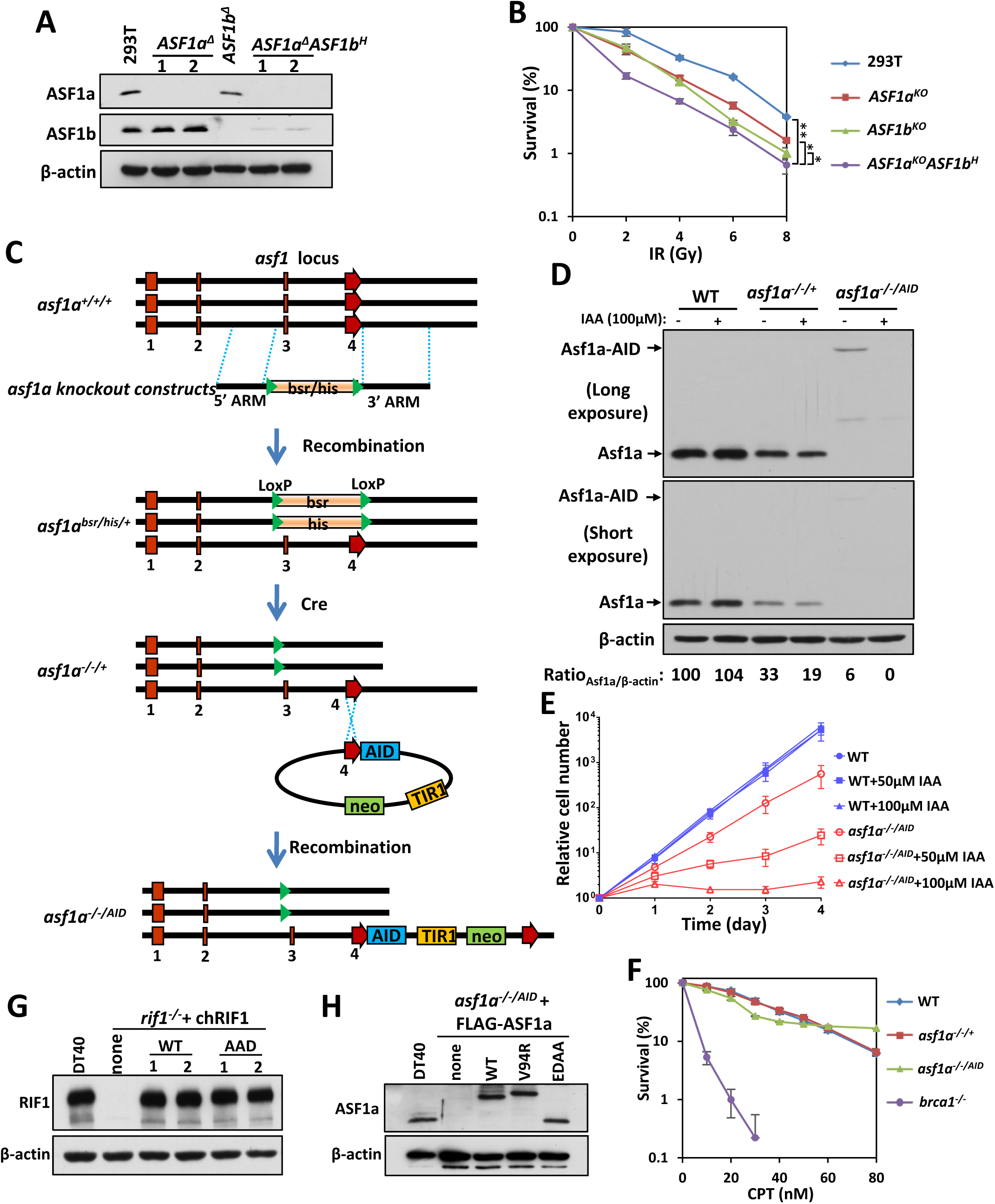
Generation of ASF1a knockout HEK293T and DT40 cells. (**A**) Immunoblots showing the expression levels of ASF1a and ASF1b in knockout cells. (**B**) IR sensitivity assay for *ASF1a^Δ^*, *ASF1b^Δ^* and *ASF1a^Δ^ASF1b^H^* HEK293T cells. The mean and s.d. of the results from two independent experiments are shown. *, p<0.05; **, p<0.01. Statistical analysis was performed using the two-tailed *t*-test. (**C**) Schematic representation of the chicken wild-type and targeted genomic DNA in the *asf1a* gene. The regions containing exons (marked by red) of these genes were replaced by histidinol or blasticidin resistance genes. The two regions between two pairs of dotted lines were used as the arms of the knock-out constructs. The third allele was fused with AID at the C-terminal. (**D**) Immunoblots showing the ASF1a level in the knockout cells with or without IAA. (**E**) Growth curves of *asf1a^-/-/AID^* cells. (**F**) CPT-sensitivity assay of *asf1a^-/-/+^* and *asf1a^-/-/AID^* cells. The mean and s.d. of the results from three independent experiments are shown. (**G, H**) Immunoblots showing RIF1 (G) and ASF1a (H) levels in their complemented cells.

**Supplemental Figure S5.**
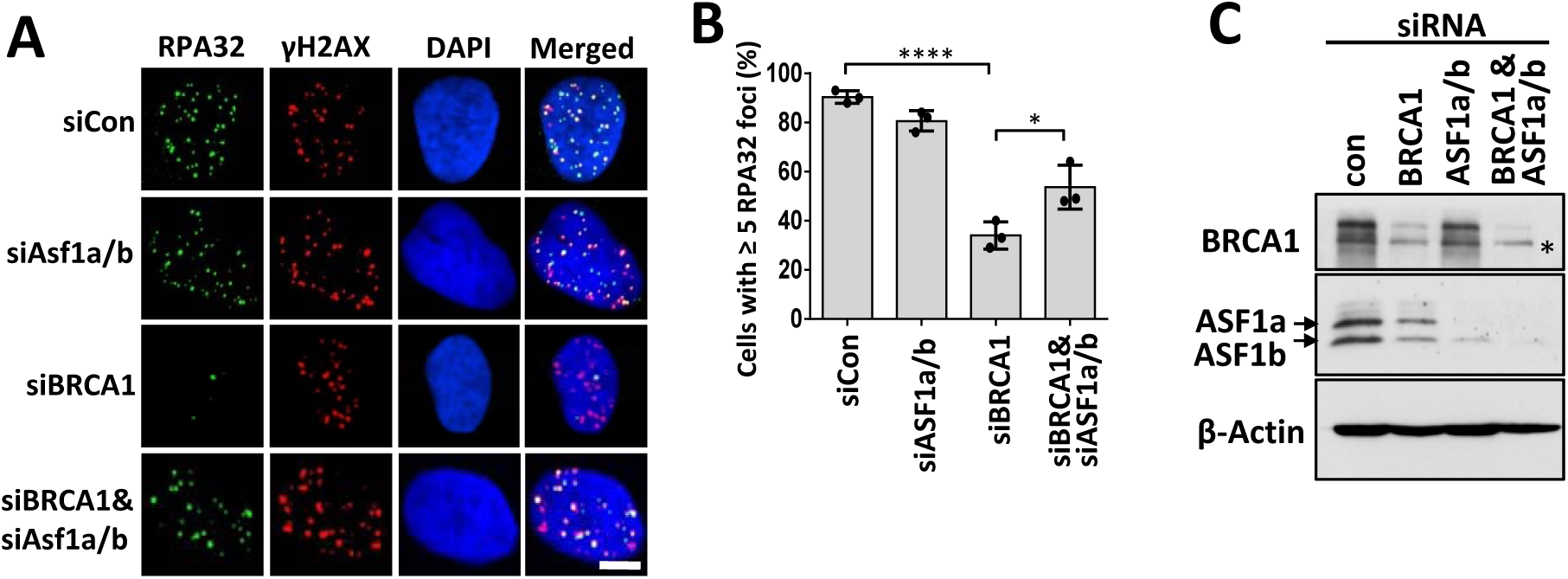
ASF1 suppresses broken end resection in BRCA1-deficient cells. (**A, B**) Immunofluorescence (A) and quantification (B) of RPA32 foci. BRCA1- or/and ASF1a/b-depleted U2OS cells were treated with 25 Gy X-ray radiation 1 h before staining. The mean and s.d. from three independent experiments are shown. Scale bar, 5 μm. ns, p>0.05; *, p<0.05; ***, p<0.001; ****, p<0.0001. Statistical analysis was performed using the two-tailed *t*-test. (**C**) Immunoblots showing the knockdown efficiency of BRCA1 and ASF1a/b in U2OS cells. Crossreactive polypeptides are indicated by an asterisk.

**Supplemental Figure S6.**
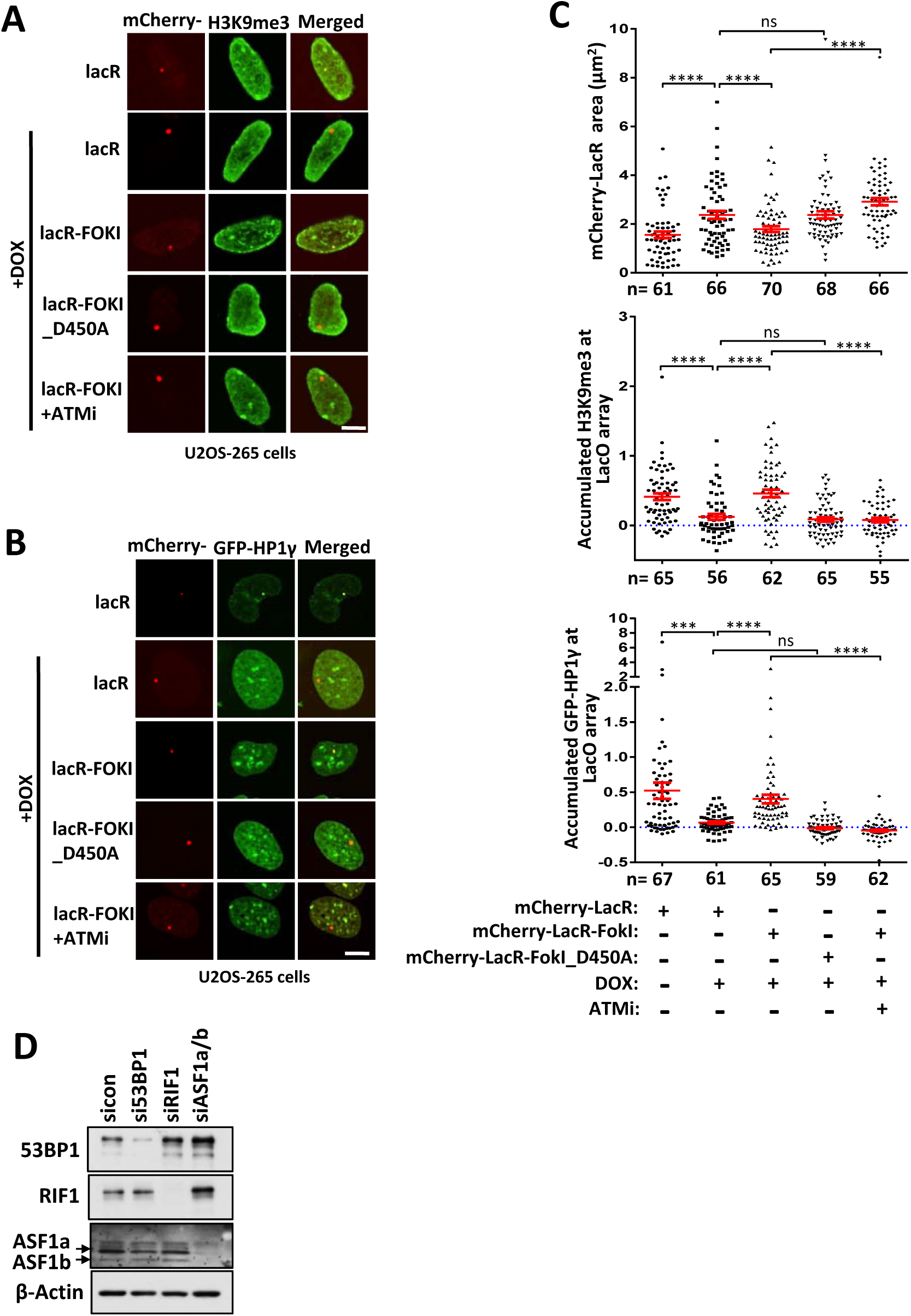
DSBs induce chromatin condensation. (A-C) H3K9me3 (A) and GFP-HP1γ (B) signals in the array and their quantification (C) in U2OS-265 cells. Cells were transfected with mCherry-LacR, mCherry-LacR-fused wild-type or D450A mutant FokI. DOX and ATMi were added 5 hr and 3 hr before imaging, respectively. The mCherry-LacR area was also quantified in (C). (**D**) Immunoblots showing the knockdown efficiency of 53BP1, RIF1 and ASF1a/b in U2OS-265 cells. Scale bar, 5 μm. ns, p>0.05; ***, p<0.001 ; ****, p<0.0001. Statistical analysis was performed using the two-tailed *t*-test.

**Supplemental Figure S7.**
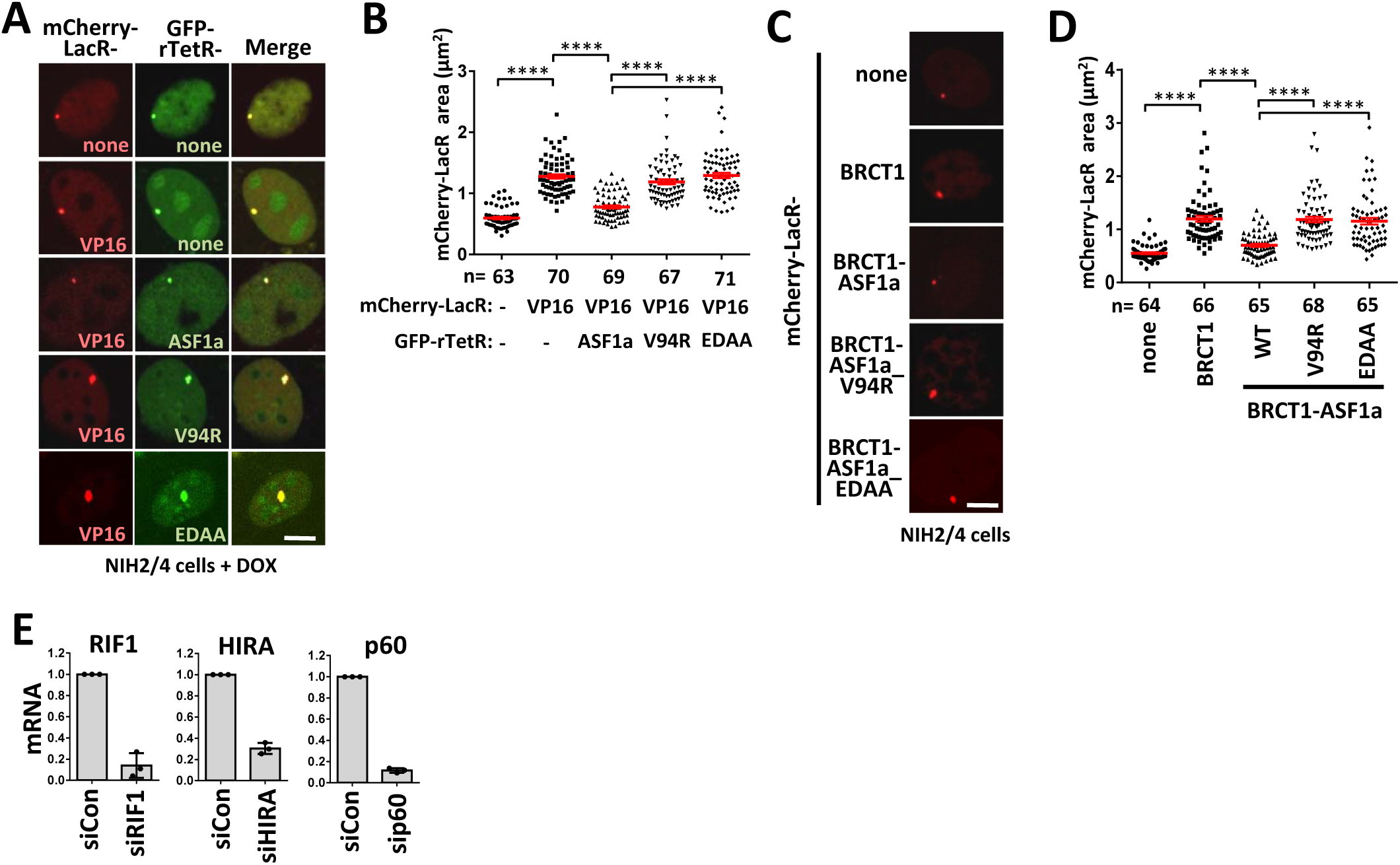
ASF1 condenses opened-chromatin under non-DSB conditions. (A-B) Images (A) and quantifications (A) of NIH2/4 cells expressing GFP-rTetR-fused ASF1a or its mutants (V94R or EDAA), and mCherry-LacR-fused with 1 μg/mL DOX. The mCherry-LacR area in the array was quantified. The error bars represent the SEM. **(C, D)** Images (C) and quantification (D) of NIH2/4 cells expressing mCherry-LacR-BRCT1-fused ASF1a or its mutants (V94R or EDAA). The error bars represent the SEM. **(E)** mRNA levels of RIF1, HIRA and CAF-1 p60 in U2OS-265 cells after siRNA knockdown. mRNA levels were measured by RT-PCR. The mean and s.d. from three independent experiments are shown. Scale bar, 5 μm; ****, p<0.0001. Statistical analysis was performed using the two-tailed *t*-test.

**Supplemental Figure S8.**
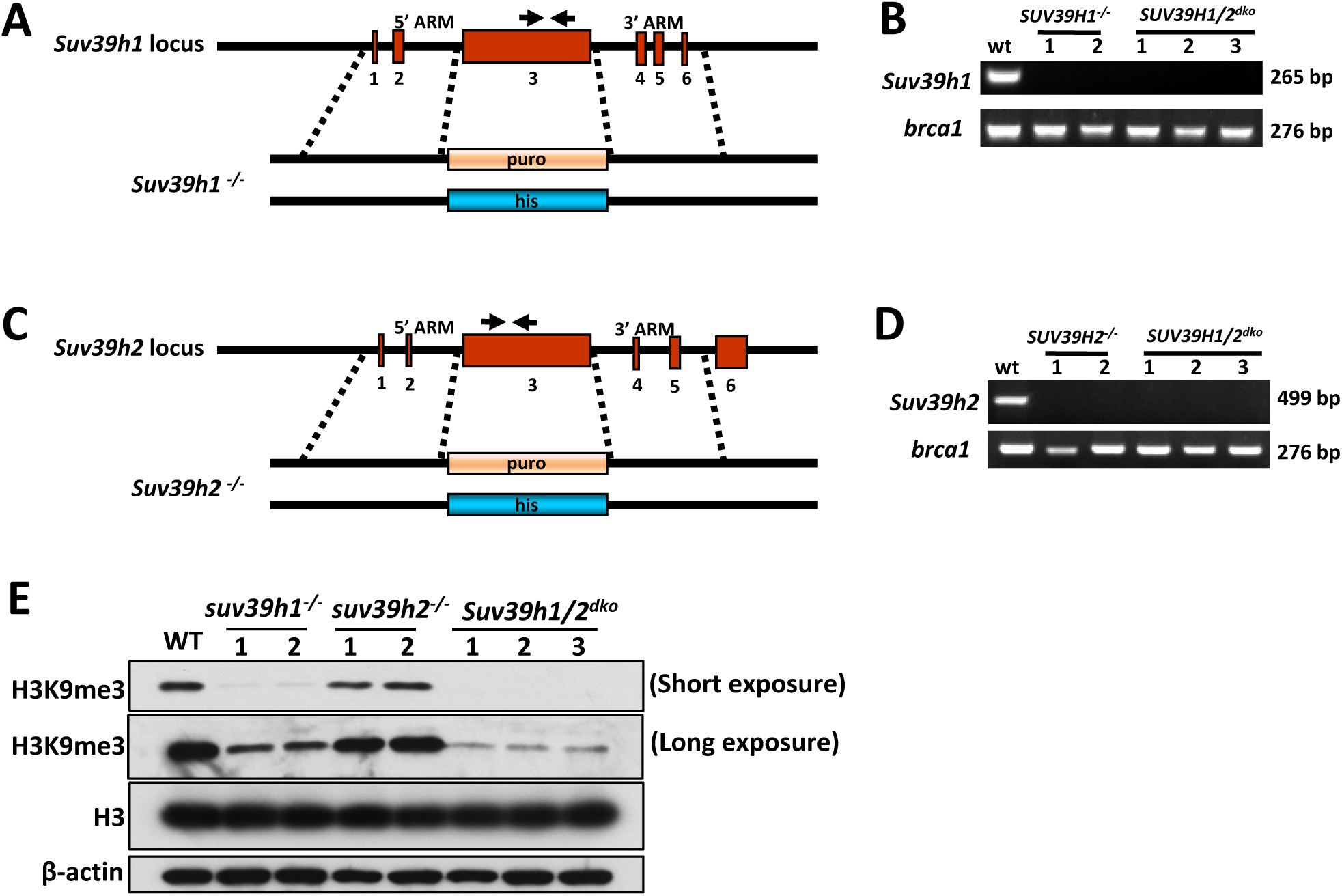
Generation of SUV39h1/2 knockout DT40 cells. (**A**, **C**) Schematic representation of the chicken wild-type and targeted genomic DNA in the *suv39h1* (A) and *suv39h2* (C) genes. Regions containing exons (marked by red) were replaced by histidinol or puromycin resistant genes. The two regions between two pairs of dotted lines were used as the arms of the knock-out constructs. Arrows indicated targeted locations of the primers for genomic PCR. (**B**, **D**) Genomic PCR analysis to show that *suv39h1* (B; primers: TTCGTCTACATCAACGAGTACAAAGT and AAGATGCAGAGGTCGTAGCGGAT) and *suv39h2* (D; primers: GGAAAGGATGGCCAGAATCTT CAAATACTTGGGAACC and CTGCAAAATGAATTGCATTCATAGATGGGCAAGCC) genes are undetectable in the respective knockout DT40 cells. *Brca1* (primers: CAATTGAGGGCCAGAGTTCTC and GAGAATCATCC ATGGGTTCACAC) was included as a positive control. (**E**) Immunoblots showing that H3K9me3 is reduced in the knockout cells.

**Supplemental Figure S9.**
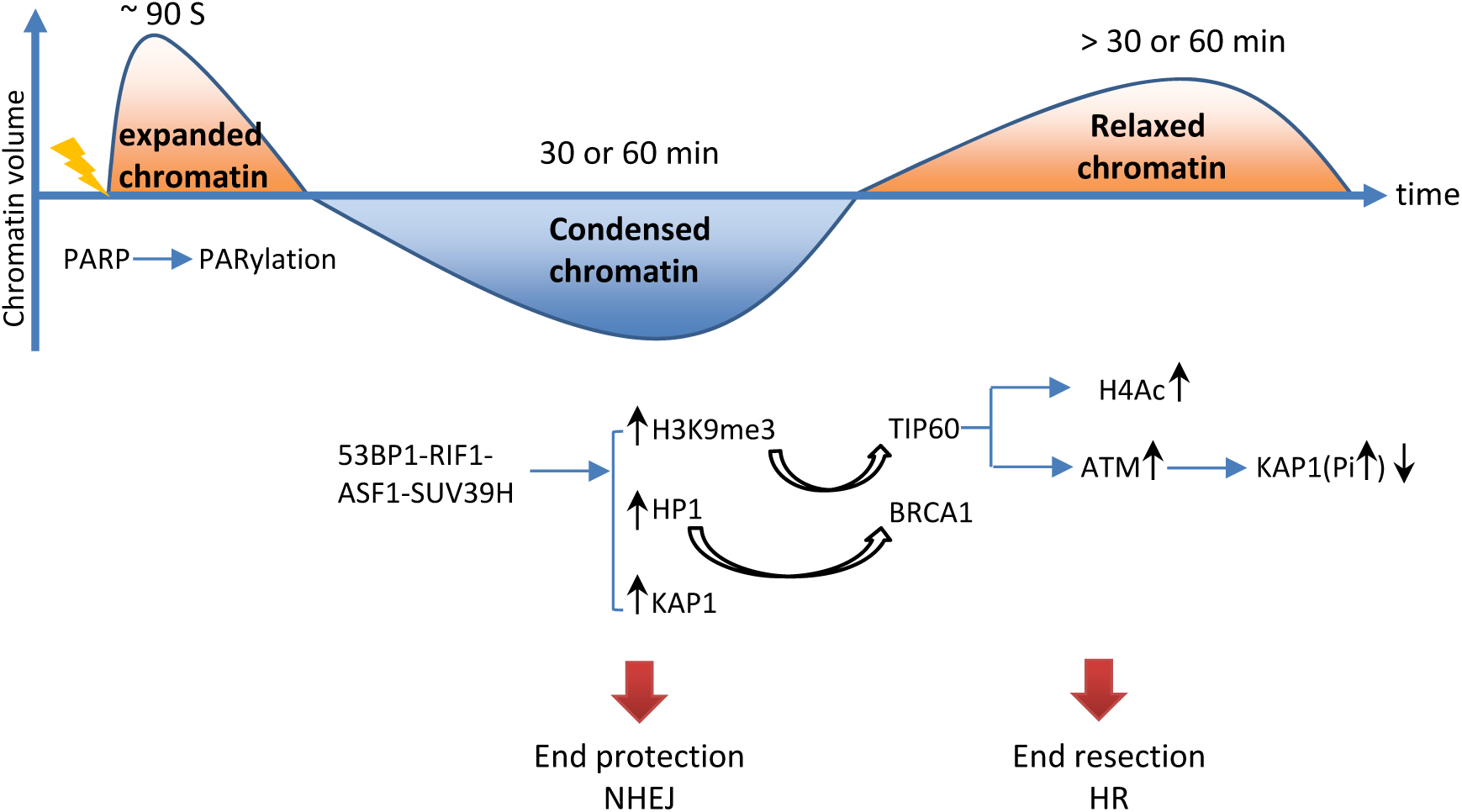
A model for chromatin dynamics during DSB repair. DSB induces a transient chromatin expansion in 90 seconds in a PARylation-dependent manner. Then, the chromatin undergoes a condensation through heterochromatinization mediated by the 53BP1-RIF1- ASF1-SUV39H axis. This condensation protects the broken ends from resection, and thus the DSBs tend to be repair by NHEJ. If rapid repair by NHEJ does not work (in 30 or 60 min), the accumulated heterochromatin marks will recruit TIP60 and BRCA1, which lead to chromatin relaxation and end resection. And then, the DSBs tend to be repaired by HR.

